# How to quantify developmental synchrony in malaria parasites

**DOI:** 10.1101/2024.02.27.582296

**Authors:** Megan A. Greischar, Nicholas J. Savill, Sarah E. Reece, Nicole Mideo

**Affiliations:** Department of Ecology & Evolutionary Biology, Cornell University, Ithaca, NY, US; Department of Ecology & Evolutionary Biology, University of Toronto, Toronto, ON, Canada; Institute of Ecology and Evolution, University of Edinburgh, Edinburgh, UK; Institute of Immunology and Infection Research, University of Edinburgh, Edinburgh, UK

**Keywords:** synchrony, model-validated methods, *Plasmodium chabaudi*, *Plasmodium falciparum*, population dynamics, developmental timing, intraerythrocytic development, Leslie matrix

## Abstract

Malaria infections represent an iconic example of developmental synchrony, where periodic fevers can result when the population of parasites develops synchronously within host red blood cells. The level of synchrony appears to vary across individual hosts and across parasite species and strains, variation that—once quantified—can illuminate the ecological and evolutionary drivers of synchrony. Yet current approaches for quantifying synchrony in parasites are either biased by population dynamics or unsuitable when population growth rates vary through time, features ubiquitous to parasite populations *in vitro* and *in vivo*. Here we develop an approach to estimate synchrony that accounts for population dynamics, including changing population growth rates, and validate it with simulated time series data encompassing a range of synchrony levels in two different host-parasite systems: malaria infections of mice and human malaria parasites *in vitro*. This new method accurately quantifies developmental synchrony from per capita growth rates using obtainable abundance data even with realistic sampling noise, without the need to sort parasites into developmental stages. Our approach enables variability in developmental schedules to be disentangled from even extreme variation in population dynamics, providing a comparative metric of developmental synchrony.

## 1 INTRODUCTION

Malaria parasites replicate in the blood of their hosts, and when this replicative cycle is synchronized, it can result in the periodic fevers classically associated with malaria infection (Kitchen, 1949; Nerlich et al., 2008). Synchrony indicates the degree to which parasites within infected red blood cells (iRBCs) show similar developmental timing across the population of iRBCs that make up an infection, and it can have consequences far beyond triggering symptoms. Since antimalarial drugs are most effective against certain parasite developmental ages (ter Kuile et al., 1993), parasite populations may be vulnerable—or resilient—to clearance by fast-acting drugs depending on the timing and synchronization of development (White et al., 1992). Theory suggests that pathogenic organisms could evolve different developmental schedules as a form of non-classical drug resistance (Neagu et al., 2018). Such evolutionary changes in synchrony could have knock-on consequences for malaria epidemiology, since synchrony is thought to influence infection severity (Touré-Ndouo et al., 2009) and the odds of onward transmission (Greischar et al., 2014; Schneider et al., 2018). Synchrony also influences capacity to accurately measure population expansion, generating spuriously large parasite multiplication rates and making it challenging to quantify how novel therapeutics (e.g., drugs, vaccines) impact parasites *in vivo* (Greischar and Childs, 2023). Anticipating synchrony and its potential to evolve requires understanding the degree to which parasites show heritable variation in their developmental schedules, but quantifying that variation has so far been stymied by methodological challenges. Existing methods are incapable of quantifying synchrony in ways that enable comparisons across genetic backgrounds and environments (Greischar et al., 2019).

The difficulty of quantifying synchrony arises in diverse systems (including insect pests, Bjørnstad et al., 2016), but addressing the challenges for malaria parasites—where data are available from artificial culture, experimental animal models, and natural infections—offers scope for making the comparisons needed to resolve the proximate and evolutionary drivers of developmental rhythms. In the context of malaria parasites, synchrony refers to the period of development and multiplication within a red blood cell (RBC)—the “intraerythrocytic development cycle” or IDC—that is completed when the RBC bursts to release merozoites, stages capable of invading RBCs. In practice, synchrony is nearly always quantified as the percentage of parasites in a particular age range (i.e., a morphologically distinct stage of the IDC, Lambros and Vanderberg 1979; Deharo et al. 1994, 1996; Reilly et al. 2007; Touré-Ndouo et al. 2009; Allen and Kirk 2010; O’Donnell et al. 2011), a metric that suffers from multiple sources of bias. Attempting to distinguish developmental age from morphology is inherently subjective (Ciuffreda et al., 2020) and the duration of a particular morphological stage can vary across genotypes (e.g., the early ‘ring’ stage in *Plasmodium falciparum*, Reilly et al., 2007).

Distinct from these practical challenges, stage percentages represent a biased estimate of synchrony because parasite age distributions vary with population growth rates (White et al., 1992; Khoury et al., 2014; Greischar et al., 2019). This bias will be far more widespread—impacting infections *in vivo* and *in vitro*—since malaria infections exhibit extreme variability in population dynamics, such that parasite numbers within an infection can vary over several orders of magnitude on the timescale of days or weeks (e.g., Miller et al., 1994; Huijben et al., 2010; Wacker et al., 2012). Robust comparisons can be made by focusing on time windows where population dynamics are similar across treatments (e.g., comparing synchrony in typical versus perturbed host feeding rhythms, Prior et al. 2018), but that approach cannot be generalized, e.g., to the case where treatments alter parasite population dynamics. *In vitro* experiments reflect this quandary: Allen and Kirk (2010) hypothesized that shaking *P. falciparum* cultures would help maintain synchrony by reducing spatial variation in nutrient availability and concomitant variability in developmental duration, and they tested that hypothesis by checking for a greater percentage of iRBCs in the early part of development in shaken versus static cultures following initial synchronization. However, shaking cultures also accelerates population expansion by reducing merozoites that are wasted invading RBCs that have already been infected, as would be frequent in static cultures (Allen and Kirk, 2010). Thus, a greater percentage of young iRBCs would be expected even if shaking did not reduce variability in developmental duration. Recent efforts to quantify synchrony have focused on using gene expression data to obtain parasite age distributions (Ciuffreda et al., 2020), but the fundamental problem remains: age distributions, no matter how well-resolved, are still influenced by population dynamics in ways that preclude broader comparison of developmental synchrony (Greischar et al., 2019).

The population dynamics that undermine the stage percentage approach also preclude the most commonly used method for quantifying synchrony that does not rely on age distribution data. In some malaria species, iRBCs in the latter half of parasite development sequester in the microvasculature of their hosts where they cannot be easily sampled (e.g., *P. falciparum*, White et al. 1992), causing periodic fluctuations in iRBC abundance where the period corresponds to the duration of the IDC. The fluctuations can be fit with a model to quantify synchrony. Those models assume log-linear expansion (i.e., a constant rate of parasite multiplication) and fit a sine wave, the amplitude of which serves as a metric for the degree of synchrony (e.g., Simpson et al. 2002, reviewed in Gnangnon et al. 2021). That approach sidesteps the bias inherent to stage percentages but cannot be applied broadly, for example, when sequestration is absent or when parasite multiplication rates change through time. Sequestration is not ubiquitous and cannot take place *in vitro* or in certain hosts (e.g., *rag1-/-* mice, Khoury et al. 2014). While parasites often multiply exponentially initially, that log-linear expansion cannot continue indefinitely as resources become exhausted and immune pressure mounts, so the latter portions of time series must be discarded prior to fitting the model. It is also not obvious how the amplitude of fitted sine waves can be translated into a biologically meaningful and readily comparable measure of developmental synchrony, especially for malaria species that do not sequester and hence do not exhibit oscillations in observable iRBC abundance.

Here we present a new method for quantifying synchrony that treats the age distribution as unobserved data and instead quantifies developmental synchrony by fitting a simple model, a Leslie matrix (Leslie, 1945), to infection time series (specifically, repeated measures of infected RBC densities). This approach estimates the time window over which a cohort of parasites within an infection complete their IDC and burst out of RBCs: a short window, relative to the cycle length, indicates a highly synchronized infection, while a long window indicates asynchrony. Since no existing approach can provide accurate estimates of synchrony from empirical data against which to compare (Greischar et al., 2019), we instead validate this approach by applying it to simulated data (i.e., where the true answer is known). We do this for two malaria species, *Plasmodium falciparum* and *P. chabaudi*, that show considerable differences in the duration of the IDC and population dynamics. Our approach can recover differences in synchrony using tractable sampling schedules, and, equally important, can detect similar levels of synchrony despite differences in IDC duration and population dynamics across species. This method relies on iRBC counts sampled multiple times per IDC rather than labor-intensive morphological staging. The focus on stage percentages has left a dearth of relevant datasets, but these data are readily obtainable whenever sequestration does not obscure iRBC abundance (e.g., *in vitro*). Unlike existing metrics, this definition of synchrony enables comparisons across species and environments even when population dynamics also vary, providing a framework for understanding the evolutionary drivers and practical consequences—including the capacity for harm and vulnerability to control—of synchrony.

## 2 METHODS

We develop an approach for measuring synchrony by fitting a model to per capita replication rates calculated from iRBC time series. We estimate synchrony from the age distribution of parasites within iRBCs, similar to past efforts (Ciuffreda et al., 2020) but with an important difference: instead of directly measuring age distributions—a composite measure of synchrony and population dynamics—we fit the initial age distribution with a model that accounts for subsequent changes in age structure caused by population dynamics. Ideally, we would apply this method to real data, comparing its performance against the current gold standard approach. However, in this case the gold standard metric—the percentage of parasites in a particular developmental window—is known to be unacceptably biased (Greischar et al., 2019), so we instead simulate data where the true level of synchrony is known (that is, specified in the model) and use those simulated data both to illustrate the bias inherent to the gold standard approach and to validate our novel approach.

We use a previously described mechanistic model (Greischar et al., 2014) to represent experimental infections of mice with the rodent malaria parasite, *P. chabaudi*, and *in vitro* cultures of human malaria parasites, *P. falciparum*. These two scenarios exhibit divergent population dynamics: *P. chabaudi* requires 24 hours for the IDC and RBCs are replenished by the host causing iRBC abundance to rebound from infection-induced decline; *P. falciparum* requires 48 hours for the IDC and replicates in an artificial culture of RBCs without RBC replenishment (in our simulations). We simulate four different levels of synchrony, identical for each of the two scenarios (Fig. 1), and sample our simulated infections on empirically feasible schedules. To test low resolution time series, we simulate ‘even’ sampling where infections are censused twice per IDC with equal time between each sample (12 hours apart for *P. chabaudi* and 24 hours apart for *P. falciparum*). We also examine ‘uneven’ sampling where more samples are taken per IDC and sampling is concentrated around peak bursting. For uneven sampling, we assume only three samples per IDC are possible for *P. chabaudi* infections *in vivo* (gathered in an eight hour window around peak bursting), whereas we simulate sampling *P. falciparum in vitro* every two hours for the 12-hour period spanning peak bursting (seven samples per IDC, as described by Reilly Ayala et al., 2010). By using distinct sampling schedules for the two species, we also test whether the new approach can cope with practical differences in the timing and frequency with which malaria species can be assessed.

**Figure 1.**
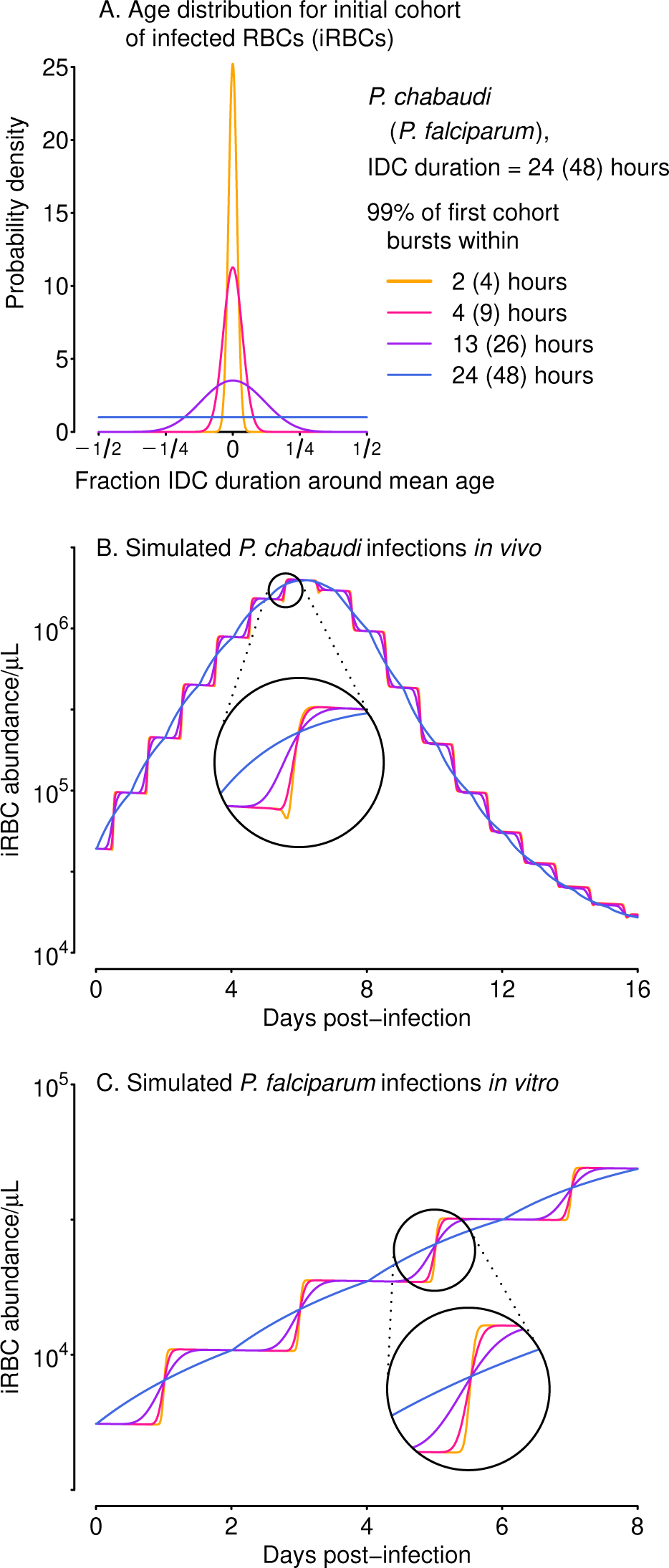
Differences in synchrony are dwarfed by population dynamics, even for unrealistically well-sampled time series. We used a range of starting age distributions for the initial cohort of iRBCs (A) and simulated iRBC abundance for *P. chabaudi* infections of mice (B) and *P. falciparum* infections *in vitro* (C), both ‘sampled’ every 15 minutes. The initial age distribution for each cohort is described by a symmetric beta distribution and is assumed to be extremely synchronous (a narrow distribution of parasite ages, *s_P_* = 500, orange), highly synchronous (*s_P_* = 100, pink), synchronous (*s_P_* = 10, purple), or asynchronous (a uniform distribution of parasites ages, *s_P_* = 1, blue). Those beta distributions translate into 99% of the initial inoculum bursting within 2, 4, 13 or 24 hours (respectively) for *P. chabaudi*, or within 4, 9, 26 or 48 hours (respectively) for *P. falciparum*.

We use simulated iRBC abundance as is and also with realistic sampling noise. We assume that iRBC abundance will be assessed via microscopy, since qPCR estimates parasite genome copies that can only be related to iRBC abundance if the level of synchrony is known (e.g., Cheesman et al., 2003). Microscopy protocols for estimating iRBC abundance via microscopy vary, so we test the consequences of either counting iRBCs until a target total number of RBCs have been observed (binomial error) or counting to a target iRBC number (negative binomial error). Details of the model, parameter values, and simulated sampling schedules and sampling noise can be found in Supplemental Methods. We confirm that simulated data generate biased estimates of synchrony using existing methods (percentage of iRBCs in the early half of development) and then test our new approach for quantifying synchrony.

### 2.1 A new metric for developmental synchrony

The iRBC age distribution—estimated while accounting for changes due to population dynamics—can be used to demarcate the time required for 99% of bursting (as in figure 1A), and dividing that quantity by the duration of the IDC can serve as a measure of developmental synchrony that can be readily compared across species:

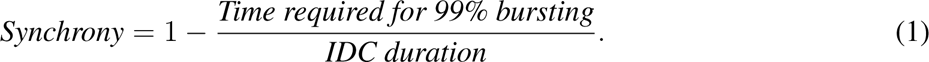

We subtract the fraction from one to arrive at a measure of synchrony that scales from zero (completely asynchronous) to one for perfectly synchronized infections, consistent with past efforts to establish a “synchronicity index” (Deharo et al., 1996). The time required for 99% bursting corresponds to the 0.005 and 0.995 quantiles of the Beta distribution given by the shape parameter *s_P_*, multiplied by the duration of the IDC. Thus, for the same shape parameter, the time required for 99% of bursting would take twice as long for *P. falciparum* compared with *P. chabaudi* (figure 1), since *P. falciparum* requires twice as long to complete its cycle (reviewed in Paul et al. 2003; Mideo et al. 2013).

We use a Beta distribution because it is flexible enough to yield age distributions ranging from uniform to narrow bell curves (figure 1). The choice of 99% of the time required for bursting can yield values comparable to the full duration of the IDC for asynchronous infections, and hence a synchrony of zero according to Eq. 1. Note that synchrony can be calculated given an age distribution (here specified by one parameter, *s_P_*) and the duration of the IDC. We test our approach on simulated data by comparing known *s_P_* values (i.e., the values used to simulate the time series) with estimated values (*s*^*_P_*) obtained by fitting a simple model (next section). We examine the simpler case of synchrony being maintained through time and later discuss how this approach could encompass changing synchrony.

### 2.2 Projecting population dynamics using Leslie matrices

We focus on one of the simplest ways of modeling the continual feedbacks between population dynamics and age distributions, a Leslie matrix (Leslie 1945; reviewed in Hastings 1997). Given information on the number of new iRBCs generated by each bursting schizont, a Leslie matrix can be used to project the population expansion (or decline) of an age-structured population, illustrated with a hypothetical example of a population with four age classes in fig. 2. In the context of malaria infections, a Leslie matrix can be used to specify how iRBCs containing parasites of different ages transition to subsequent developmental ages or burst to generate new iRBCs (fig. 2B).

**Figure 2.**
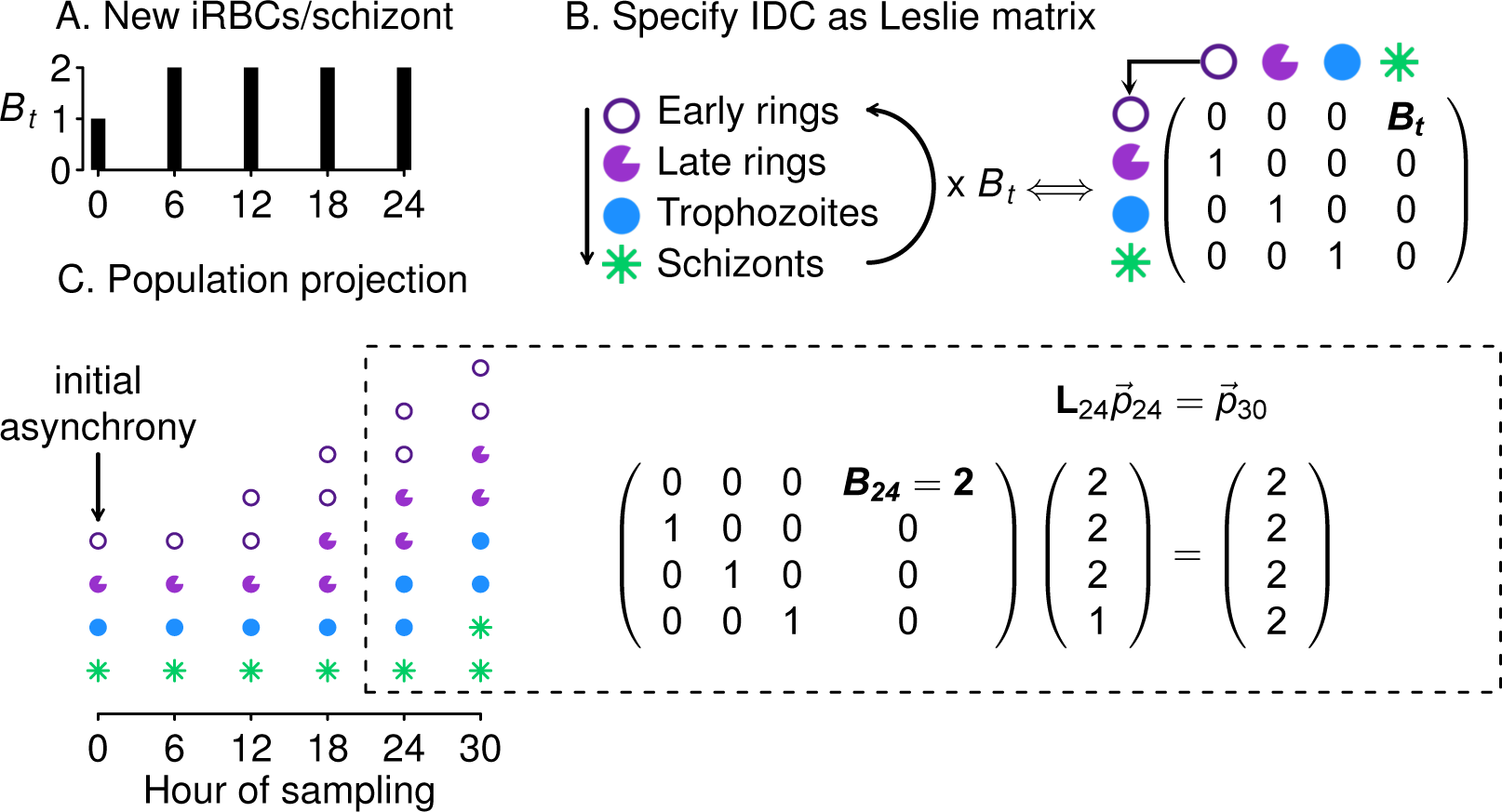
Leslie matrices project how age distributions change in response to changing population growth. Panel A shows a hypothetical example of changing numbers of new iRBCs produced per schizont (*B_t_*) through time. In panel B, progression through development is shown as a life cycle diagram (left) and, equivalently, as a Leslie matrix (right). Leslie matrices—one for each *B_t_* value—are used to project how age distributions change through time (C), as in the example calculation shown at right. The Leslie matrix is denoted **L** while *p_* indicates the abundance of iRBCs in each developmental class.

We assume that the iRBC population is initially distributed across *n* developmental age classes, where 1*/n* indicates the fraction of the IDC spent within each class (e.g., if samples were taken, at minimum, 4 hours apart for a species with an IDC lasting 24 hours, then *n* = 24*/*4 = 6, and each age class lasts 4 hours). When iRBCs are sampled at evenly spaced intervals throughout the developmental cycle, *n* is equal to *k*, the number of samples per cycle (e.g., in fig. 2, *n* = *k* = 4). If sampling is uneven, *n* is equal to the duration of the IDC divided by the minimum time between samples. For example, if sampling occured three times per cycle (*k* = 3) so that iRBCs were censused four hours prior to peak bursting, at peak bursting, and four hours after peak bursting, we treat that situation as though iRBCs were counted every four hours (with missing counts for three of the six samples), so that again *n* = 24*/*4 = 6.

For a population with *n* age classes, the Leslie matrix (**L**) is

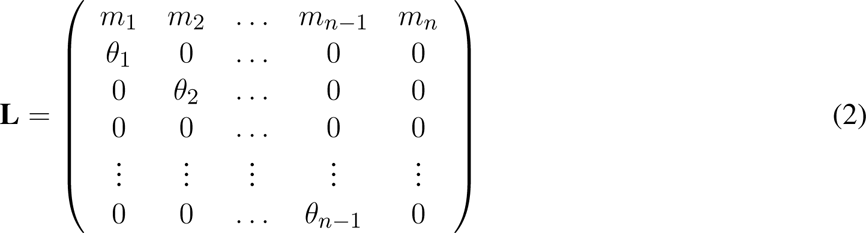

where *m* values indicate the fecundity of each age class and *θ* values give the probability of individuals in a particular age class surviving to mature into the subsequent age class in the next time point. We assumed that each age class is certain to survive and mature into the subsequent age class at the next time step (*θ*_1_ = *θ*_2_ = *θ*_3_ = *. . .* = *θ_n−_*_1_ = 1). In the simulated data, approximately 2% of iRBCs will not survive development due to background mortality of RBCs (*µ*, see Tables S1 and S2). If we are nonetheless able to recover the true duration of bursting, it would suggest the approach is robust to at least small deviations from the assumption of 100% survival of iRBCs during the IDC. Further, background RBC mortality is likely to be the dominant force removing iRBCs in the acute portion of infection, where RBC limitation can largely explain dynamics (Mideo et al., 2008) and immune clearance is thought to be minimal (White et al., 1992). Immunity will certainly be minimal *in vitro*. Note that immune clearance of merozoites would be accounted for through the calculation of parasite multiplication rates (PMRs), the fold-increase in iRBC abundance over each IDC represented in the time series (defined in Table 1). However, when immune clearance of developing iRBCs is substantial, the assumption of 100% iRBC survival through each IDC may need to be revisited.

**Table 1.** Key terms and definitions used in our metric of synchrony, as applied to malaria parasites.

| Term | Description | Notes |
| --- | --- | --- |
| $iRBC_t$ | Infected RBCs/μL blood at sampling time $t$ | Not equivalent to parasite genome copies, the output of qPCR |
| $s_P$ | Shape parameter for symmetric Beta distribution | Used to specify age distribution of iRBC initial cohort |
| $R_e$ | Per capita replication rate (fold change from one time point to the next) | $R_{e,t} = \frac{iRBC_{t+1}}{iRBC_t}$ |
| $n$ | Number of developmental age classes tracked in the Leslie matrix | |
| $k$ | Samples per cycle | With even sampling, $k = n$; for uneven sampling, $k < n$ |
| $PMR$ | Parasite multiplication rate (fold change in iRBC abundance over one IDC) | $PMR_t = \frac{iRBC_{t+k}}{iRBC_t}$ |

Since only the final age class of malaria parasites are capable of bursting out to generate new iRBCs, *m*_1_ = *m*_2_ = *m*_3_ = *. . .* = *m_n−_*_1_ = 0 and *m_n_* = *B_t_*, where *B_t_* is the number of new rings generated by each iRBC completing development (fig. 2A). Note that *B_t_* is not equivalent to the burst size, the average number of merozoites emerging from each bursting schizont, because some emerging merozoites will fail to successfully invade. Once *B_t_* values are obtained by interpolating between PMR values (see below), the Leslie matrix can be parameterized for each time point at which population projections are needed. Given a vector 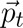 to indicate the numbers of iRBCs in each age class at time *t*,

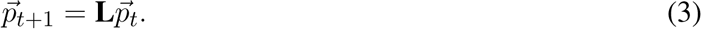

The number of iRBCs in each age class is then summed to obtain the projected total number of iRBCs at each time point. Projecting the total iRBC abundance enables calculation of the predicted per capita growth rate, *R*^^^*_e_* for each time point, where

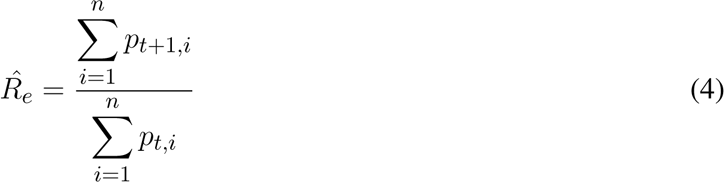

and the *i* subscript indicates the developmental age class.

The initial number of iRBCs in each developmental age class, denoted by the vector 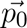, can be specified with a symmetric beta distribution, which has the useful property of ranging from uniform (i.e., completely asynchronous, when the shape parameter, *s_P_*, is equal to one) to increasingly narrow bell-shaped curves with increasing *s_P_* values (figure 1A). For a symmetric beta distribution, the mean falls at 0.5 by default— equivalent to the median parasite age falling exactly halfway through the IDC—but that need not be the case for real data. We therefore fit an ‘offset’ parameter, constrained to vary between zero and one, to allow the mean of the beta distribution (that is, the median parasite age) to deviate from 0.5. We use the cumulative density function of the beta distribution to obtain the initial stage distribution, 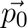, for a given number of stage compartments (*n*). The initial *I*_0_ iRBCs are sorted into *n* stages according to the beta distribution specified by a given shape parameter, *s_P_* (the quantity to be fitted). The initial number of iRBCs is arbitrary, since we are fitting per capita replication rates (rather than iRBC abundance), so we set *I*_0_ = 100 to avoid errors of precision that might occur with lower iRBC counts.

### 2.3 Estimating developmental synchrony

We describe the algorithm for estimating synchrony using Eq. 1 (illustrated in fig. 3):

**Figure 3.**
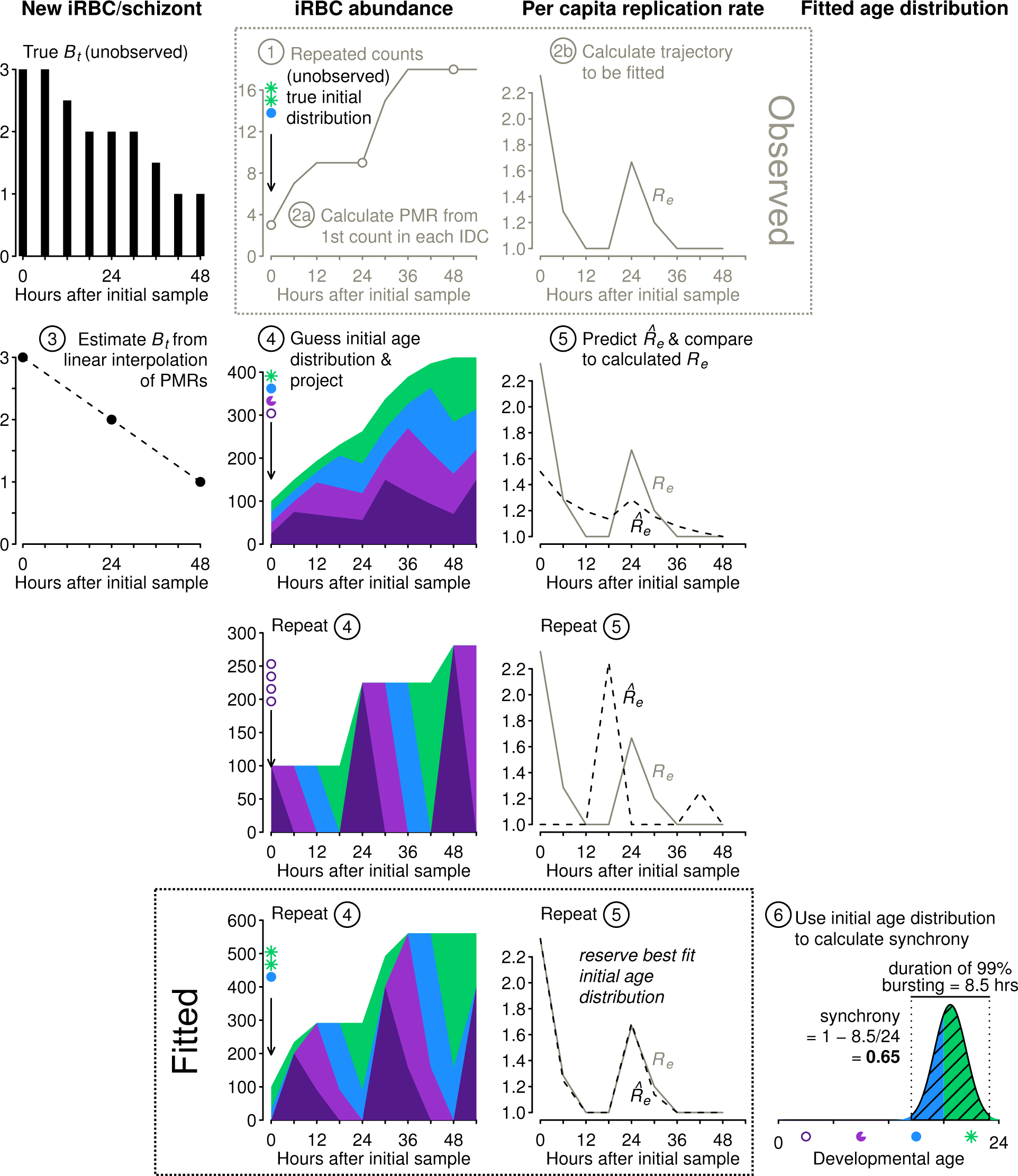
The new approach estimates synchrony while accounting for population dynamics. Example iRBC count data were obtained (Step 1) using the initial age distribution shown and projecting forward using the *B_t_* values shown at left (black, see fig. 2). We calculate PMR values (Step 2a) and per capita replication rates (*R_e_*, Step 2b), the trajectory to be fitted. We estimate *B_t_* values by linear interpolation of PMRs (Step 3). Using estimated *B_t_* values, we guess an initial age distribution and project population dynamics (Step 4) and calculate predicted *R*^^^*_e_* values to compare with observed *R_e_* values (Step 5). We repeat steps 4 and 5 and retain the best fit initial distribution to calculate synchrony using Eq. 1 (Step 6). To conserve space, y-axis labels are given at the top of each column of panels.

*Step 1*. Obtain time series of iRBC abundance, sampled multiple times per IDC. Age distribution data are not required.

*Step 2*. (a) We first calculate the parasite multiplication rates (PMRs) and then (b) calculate the per capita replication rates (*R_e_*, Table 1).

*Step 3.* By interpolating between the observed PMRs, we obtain estimated *B_t_* values needed to project population dynamics for an age-structured population (see fig. 2).

The subsequent steps are carried out numerous times, using an optimization algorithm (details follow) to determine the starting age distribution that yields predicted *R*^^^*_e_* values that best match the observed *R_e_* values:

*Step 4.* We project iRBC abundance for a large number of starting age distributions 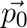 using a Leslie matrix (see example in fig. 2). The starting age distribution is specified as a symmetric Beta distribution with a shape parameter (*s*^*_P_*) that—once fitted—will be used to calculate the time required for 99% bursting and the level of synchrony according to Eq. 1. Though not used to calculate synchrony, we also fit an offset parameter to allow the median parasite age to deviate from the default of half the IDC duration.

*Step 5.* Each projected iRBC abundance is subsampled to obtain values at the same times reported in the actual data. That is, any time points that are missing in the observations are dropped prior to calculating predicted per capita replications rates (*R*^^^*_e_*) for comparison with observed per capita replication rates (*R_e_*).

*Step 6.* Identify the initial age distribution associated with the best fitting *R*^^^*_e_*. Determine the time required for 99% bursting in that initial age distribution by subtracting the 0.5% quantile from the 99.5% quantile of the Beta distribution defined by the best fit *s*^*_P_* . Calculate synchrony according to Eq. 1.

### 2.4 Numerical optimization and estimates of synchrony

We use the Nelder-Mead algorithm to locate the value of *s_P_* (and the offset parameter) that minimizes the sum of the absolute error between the observed and predicted per capita replication rates (*R_e_* and *R*^^^*_e_*, respectively). The machinery to perform the optimization is readily available from **R** (R Core Team 2020, see Supplemental methods for details and annotated code). For each simulated time series, we run the optimization 1000 times with randomly chosen starting parameters to increase the chances of arriving at a globally optimal best fit. We then retain the best fit *s_P_* and offset parameter from those 1000 optimizations (i.e., the values corresponding to the fit yielding the lowest sum absolute error) and use it to calculate the time for 99% bursting and then synchrony according to Eq. 1.

## 3 RESULTS

### Stage percentages represent biased synchrony estimates

Our simulated data can be used to show the problems associated with the common approach of quantifying synchrony as the percentage of iRBCs in a particular life stage (reviewed in Greischar et al. 2019). We split the IDC into an early and late phase of equal duration, where the early phase represents the morphologically distinguishable “ring” stage that occupies roughly the first half of intraerythrocytic development (Cheesman et al., 2003; Reilly et al., 2007). An even stage distribution—here 50% rings—would correspond to asynchrony in a population that is neither expanding or contracting, while 100% rings is typically thought to indicate perfect synchrony (Greischar et al., 2019). Though the duration of the ring stage varies across *P. falciparum* strains (Reilly et al., 2007) and presumably also across *Plasmodium* species, we make the generous assumption that the duration of the ring stage is known in advance and always occupies exactly half the IDC. We project the percentage rings at each sampled time point ±10% standard error (fig. 4, full dynamics shown in fig. S1) in line with reports from experimental rodent malaria infections in the absence of sequestration (Khoury et al., 2014). Depending on the sampling schedule, the percent rings can overlap with the asynchronous expectation for highly synchronized simulations or appear “semi-synchronized” (a bare majority of parasites in the ring stage, e.g., ~64%, Khoury et al. 2017). Note that simulations of *P. falciparum in vitro* yield higher percent rings on average, especially with poor synchrony (figure 4C, D), since that population is only expanding, in contrast to the simulations of *P. chabaudi*, which expand and then decline (figure 1B, C). The age distribution assessed at a single time point will frequently fail to detect even substantial rank order differences in synchrony, but synchrony is frequently assessed from a single time point (e.g., Touré-Ndouo et al. 2009; Ciuffreda et al. 2020).

**Figure 4.**
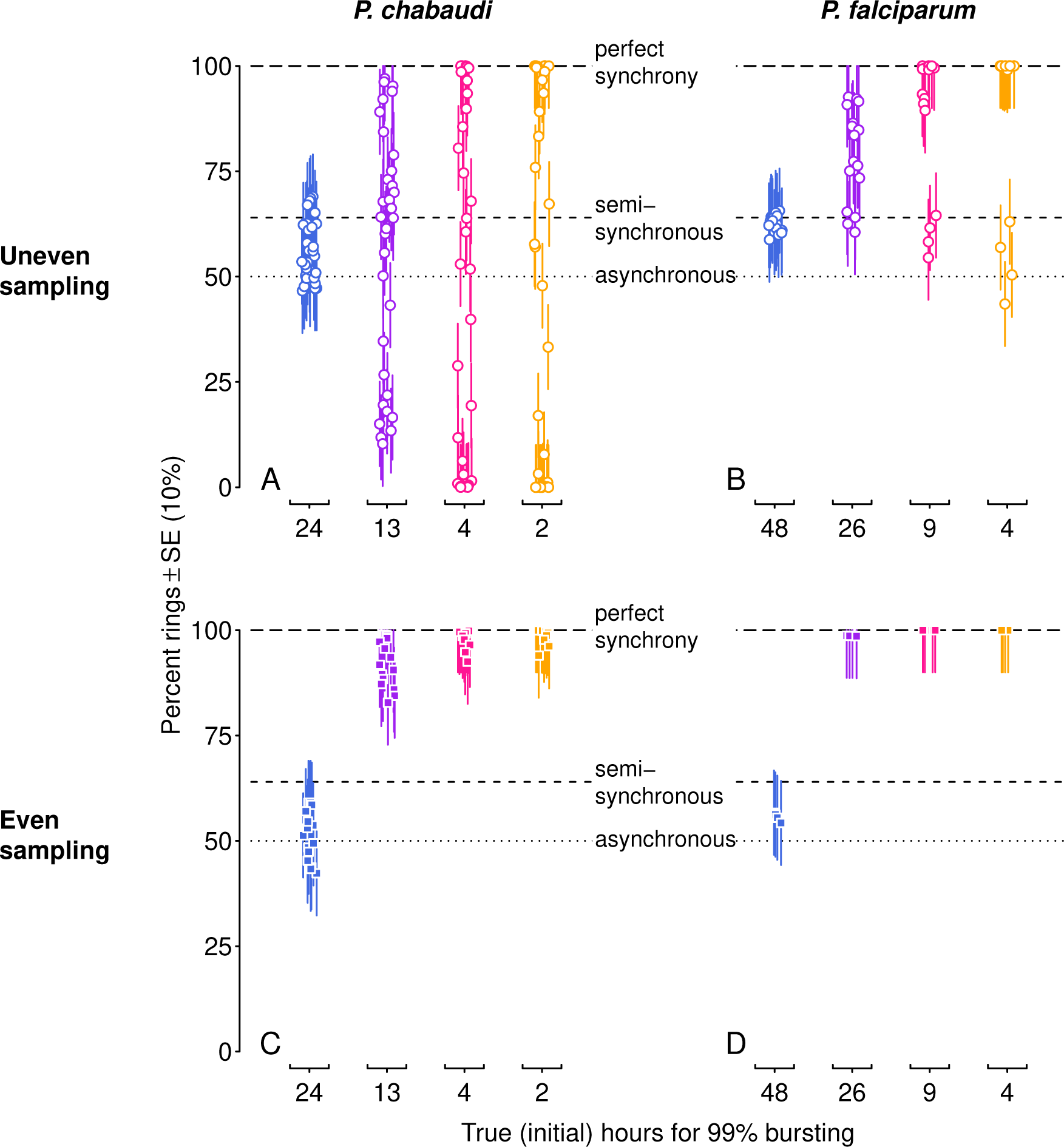
Stage percentages vary with population dynamics, making it difficult to accurately assess and compare synchrony when estimated in this way. The percentage of rings in simulated infections is shown for time points when rings would be expected (i.e., in the first half of each IDC). Ring percentages from unevenly ‘sampled’ infections (open circles in A, B) show percentages that deviate substantially from expected values, while even sampling (closed squares in C, D) can only distinguish between asynchrony and some degree of synchrony. Colors and x-axis indicate the true duration of bursting and hence initial level of synchrony as in figure 1A. *P. chabaudi* infections (A, C) were simulated over 16 IDCs, while *P. falciparum* simulations represent 4 IDCs. Vertical line segments indicate ±10%, roughly the standard error reported for stage percentage data where sequestration is not occuring (as in *rag1^−/−^* mice, Khoury et al. 2014). Horizontal lines indicate expected values for perfect synchrony (100%, long dash), asynchrony (50% rings in a static population, dotted line), or a semi-synchronous population (64% rings, following Khoury et al. 2017).

### The new approach recovers true differences in synchrony

Differences in synchrony can be difficult to discern from iRBC trajectories due to large changes in abundance that can occur in a matter of days, even with extremely high resolution time series in the absence of sampling noise (figure 1B, C). Per capita replication rates make distinct levels of synchrony much clearer (*R_e_*, figure S2A, C). However, per capita replication rates vary with the sampling schedule, and sparse sampling reduces the ability to visually detect differences in synchrony (e.g., twice per IDC, figure S2B, D). We first examined the fitted per capita replication rates compared to simulated *P. chabaudi* and *P. falciparum* values assuming no sampling noise. The simple fitted model (colors as in figure 1) yields predicted *R*^^^*_e_* values that are very close to the *R_e_* values from simulated data (figs. S3, S4).

Model fitted estimates represent the best fit from 1000 randomly chosen starting values, so we re-ran fits 20 times on simulated time series, without sampling noise, to estimate variability in the fitting process itself. When sampling was uneven, the fitted values did not vary, but even sampling twice per IDC yielded a range of estimates for the duration of bursting and hence synchrony (closed squares in figure 5). For even sampling only, multiple combinations of median parasite age (fitted with the offset parameter) and duration of bursting yielded equally good fits (figure S5). This issue—nonidentifiability—was foreshadowed by the extremely similar per capita replication rates (*R_e_* values) for all but the asynchronous simulations (figure S4). In contrast, *R_e_* trajectories are distinct for each level of synchronization with uneven sampling (figure S3) and identifiability is not an issue. We chose the timing of uneven sampling so that more samples were taken around the median time of bursting for each cohort, since that will yield the largest changes in iRBC abundance and hence the largest *R_e_* values. If the time of median bursting is known, clustering samples around that time is likely to make it easier to detect differences in synchrony. However, if the timing of bursting is not known, uneven sampling would still be useful to avoid nonidentifiability between bursting duration and median parasite age.

**Figure 5.**
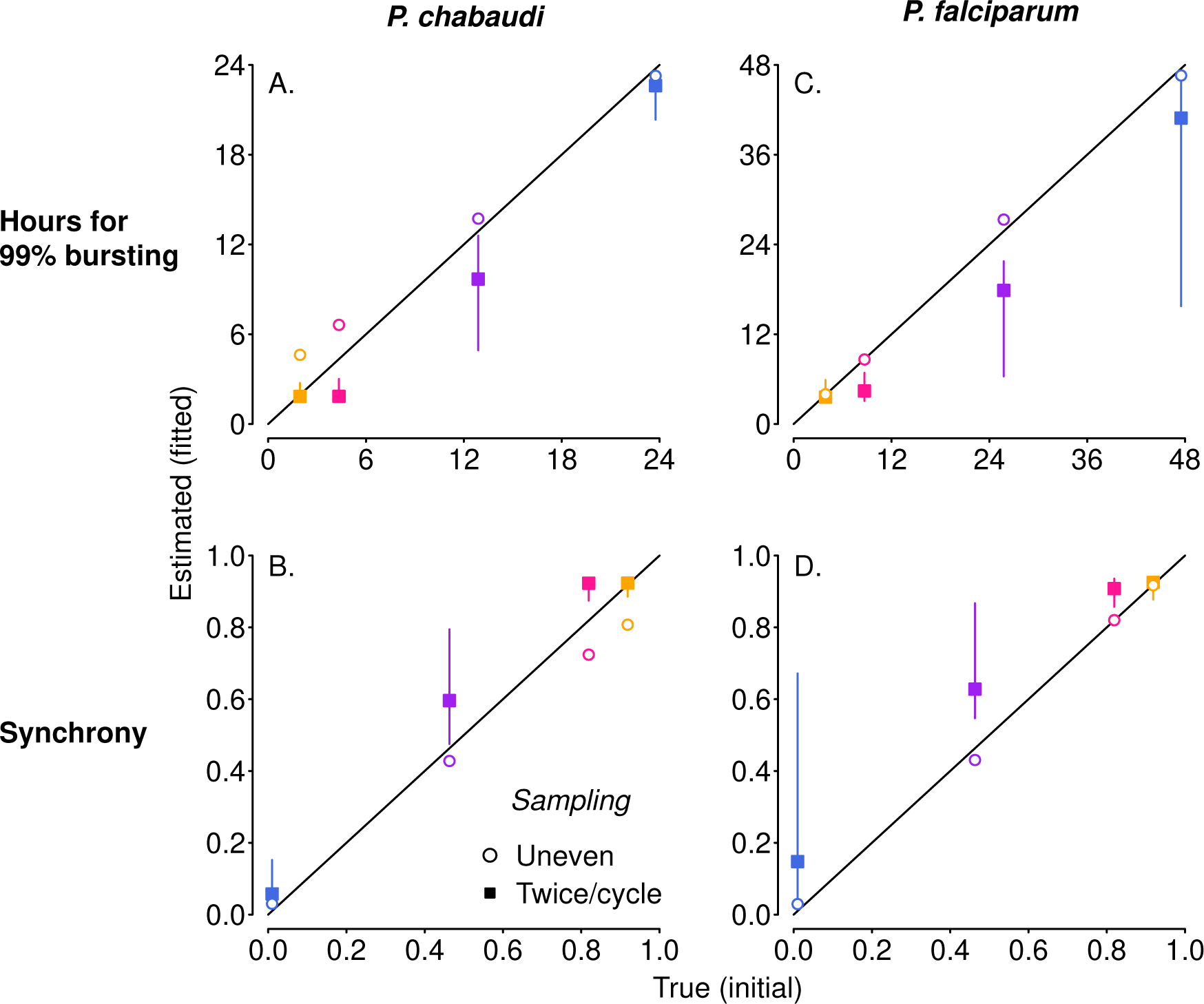
The new approach estimates the duration of 99% bursting (A and C) and synchrony (B and D) very close to the true values (i.e., the values derived from the Beta distribution used to simulate the “observed” time series), despite differences in IDC length and population dynamics. Fits to simulated *P. chabaudi* (*P. falciparum in vitro*) infections are shown in panels A and B (C and D). Open circles denote uneven sampling intervals while closed squares refer to sampling performed at even intervals twice per IDC. Vertical segments indicate the 95% high density region of predicted values; only evenly sampled time series showed any variation in the predicted values. True initial synchrony and duration of bursting is indicated with colors (as in figure 1) and by position on the x-axis. Note that in addition to better identifiability, fitting to unevenly sampled data provides more accurate estimates of true synchrony, accounting for modest loss of synchrony over simulated infections (figure S7).

Focusing on the fits to noiseless data, unevenly sampled, we obtain estimates of bursting duration and synchrony extremely close to their true initial values (open circles, figure 5). The fits to simulated *P. chabaudi* infections appear to modestly overestimate the duration of bursting, and hence underestimate the true level of synchrony (figure 5A, C), but that does not represent a problem with the method itself. Rather, these estimates are detecting a modest decline in initial synchrony over the 16 IDCs simulated, due to very slight differences in IDCs within each cohort arising from the exponentially-distributed waiting times in the merozoite stage assumed in the underlying model. High levels of initial synchrony result in discrete pulses of merozoite abundance that become wider through time as synchrony declines (figure S6). The *P. falciparum* fits are to shorter time series—4 IDCs—and the synchrony estimates are therefore nearly identical to their initial true values (figure 5B, D). When we recalculate synchrony using the mean hours for 99% bursting calculated from simulated merozoite abundance (details in supplemental code), we find that the new method returns estimates nearly identical to the true value (figure S7). Despite large differences in iRBC dynamics (figure 1B vs. C), the estimates of the duration of bursting (relative to IDC length) and synchrony are extremely similar for *P. chabaudi* and *P. falciparum* (figures 5, S7). Thus, this approach yields synchrony estimates that account for—and can therefore be uncoupled from—the population dynamics that render existing methods ineffective.

We then tested the new approach on the same underlying time series with simulated sampling noise assuming percent parasitemia was subject to a binomial or negative binomial distribution. For simulated *P. chabaudi* infections, fitted estimates of synchrony matched well with their true initial values on average, but binomial error in percent parasitemia yielded a wide range of fitted values that could sometimes suggest the incorrect rank order of synchrony (figure 6A). For negative binomial error, estimated synchrony is clustered tightly around the true value, though the highest levels of synchronization may be difficult to distinguish from one another (figure 6B). Initial fits to noisy *P. falciparum* time series using the same target counts as for *P. chabaudi* (500 for binomial noise and 100 for negative binomial) gave an unacceptably wide range of estimates and poor match to true values (figure S8). Fits to noisy *P. falciparum* time series show a wider spread of estimated values, since parameter values were chosen to keep percent parasitemia below approx. 10% (Table S2). Using larger, but still plausible, target counts (2000 and 400, respectively), we find that average fitted values can deviate from the true rank order when error is binomially distributed (figure 6C), but the true rank order can usually be recovered when error is negative binomially distributed (figure 6D). Noisy asynchronous time series present the biggest challenge to accurate estimation of synchrony, since the ratio of signal to noise is lowest. That is, oscillations in per capita replication rates (*R_e_* values) are minimal (panels D and H in figures S3, S4) so that sampling noise can easily masquerade as a higher level of synchrony. Taken together, these fits suggest that sampling to a target iRBC count should make it easier to distinguish the true level of synchrony, and that greater effort (e.g., higher target counts) will be needed for *P. falciparum* infections or other species where percent parasitemia tends towards lower values.

**Figure 6.**
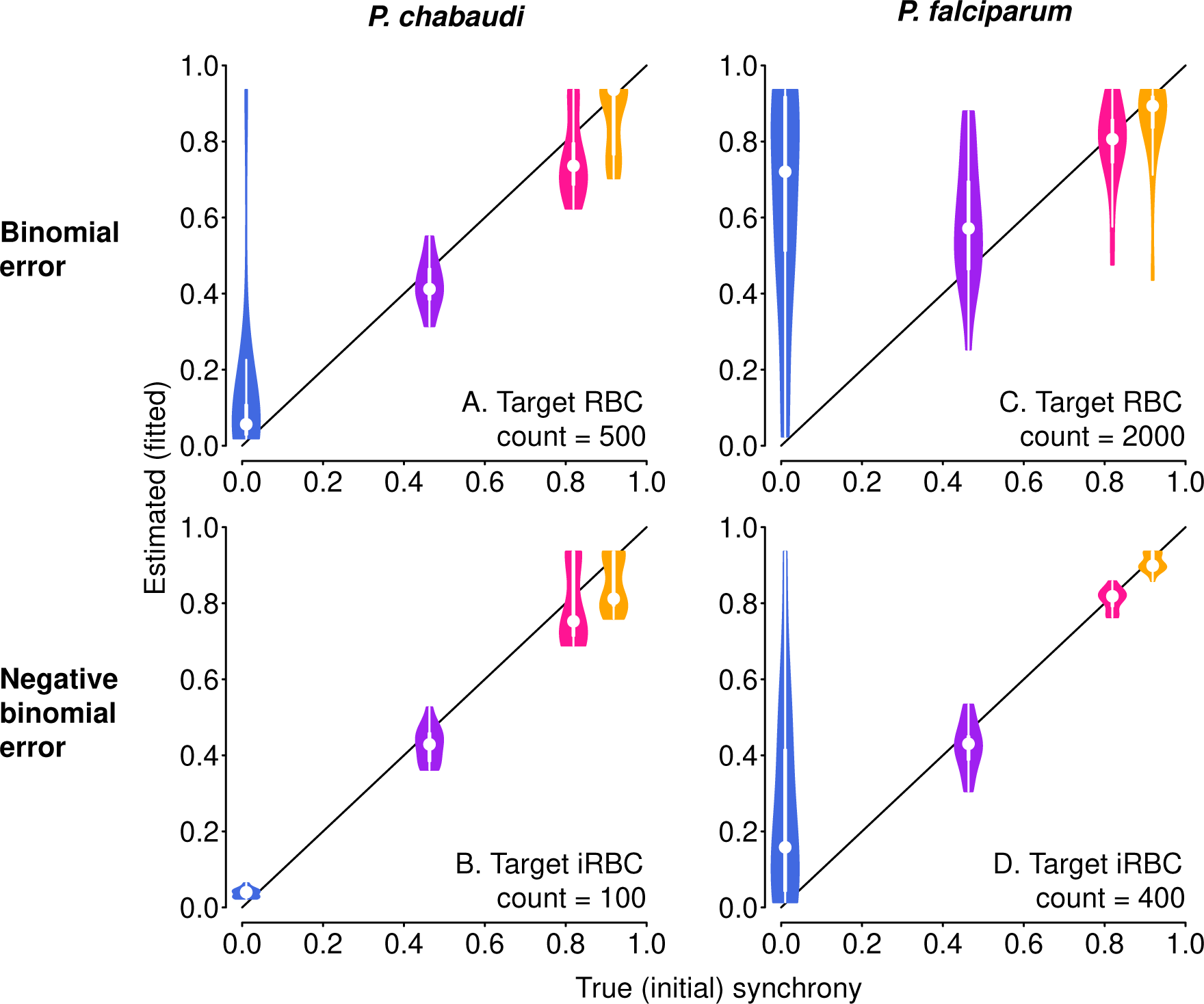
With sampling error, it is still possible to discern differences in synchrony, depending on the error distribution. Violin plots showing the distribution of 20 fits to time series with simulated sampling noise, with a binomial (A, C) or negative binomial (B, D) distribution. When error is binomially distributed (A, B), synchrony differences can be obscured, with asynchronous infections often appearing spuriously synchronous. Synchrony estimates are better behaved when sampling noise follows a negative binomial distribution (B, D). *P. falciparum* tends toward much lower percent parasitemia, especially *in vitro* (Reilly et al., 2007), so larger target RBC (iRBC) counts were used for panels C and D, following reported experimental protocols (see details in Methods section 4). For corresponding plots showing the estimated duration of bursting, see figure S9.

## 4 DISCUSSION

Developmental synchrony has the potential to alter malaria parasite fitness by influencing replication rates, drug efficacy, and transmission success (Hawking et al. 1968; Hawking 1970; White et al. 1992; Greischar et al. 2014; Schneider et al. 2018; O’Donnell et al. 2022; Owolabi et al. 2021, reviewed in Mideo et al. 2013; Greischar et al. 2019; Prior et al. 2020). Further, parasites at different developmental ages vary in their susceptibility to host defenses (e.g., *γδ* T cells, Costa et al. 2011) and antimalarial drugs (Yayon et al., 1983), including the current front-line drug, artemisinin (ter Kuile et al., 1993; Owolabi et al., 2021). The ability to account for underlying differences in population dynamics is central to comparing synchrony across environments, populations and species. Intuitive metrics (e.g., the percentage of iRBCs in a particular developmental stage) fail to account for population dynamics and yield biased estimates of synchrony (Greischar et al., 2019), except where population dynamics do not vary across treatments (Prior et al., 2018). So far the only method that sidesteps the bias generated by population dynamics is fitting a dynamical model to data (White et al. 1992; Simpson et al. 2002, reviewed in Greischar et al. 2019), but these models assume constant population growth rates, preventing comparisons within and across infections. We developed a new approach, validated against simulated data, that can distinguish the level of synchrony while accounting for considerable variation in population growth rates.

The Leslie matrix model presented here describes the dynamics for a population that begins at a particular level of synchrony and whose age distributions are subsequently unperturbed except by replication. That model is nonetheless able to return the correct average level of synchrony when the underlying dynamics deviate from the assumption of fixed IDC duration and synchrony changes through time (figure S7). As is, these methods could be applied whenever the goal is to detect average differences in synchrony, e.g., the problem of determining which treatments maintain higher levels of synchrony (such as shaking *in vitro P. falciparum* cultures to maintain uniform distribution of nutrients, Allen and Kirk 2010). When synchrony is hypothesized to be changing through time, whether due to processes internal or external to the organisms in question, the fitted Leslie matrix model represents the null expectation, serving as a starting point to compare against more complicated scenarios. For example, this approach could also be extended to quantifying how synchrony changes over the course of infection by fitting the model to portions of the time series. Each time window would then have a different fitted shape parameter *s*^*_P_*, and by extension, a different estimated duration of bursting and level of synchrony according to Eq. 1. If splitting the time series into multiple windows results in a better fit compared to fitting a single shape parameter for the entire infection—reducing the sum absolute error enough to justify the added complexity of the fitted model—that would suggest that the level of synchrony is changing through time.

Our results have implications for the type of data and the frequency and duration of sampling needed to quantify synchrony in other systems, pathogenic and free-living. Crucially, age distribution data are not required, only estimates of total population size sampled multiple times per generation. Sampling at even intervals through a generation is neither necessary nor desirable, since that makes it difficult to estimate synchrony separately from median age. We show that it is possible to quantify synchrony from as little as four generations of data, and estimating synchrony from even shorter time series may be feasible depending on the error distribution of the count data. Quantifying the error distribution requires information on the mean and standard deviation of population size from replicate samples (see Huijben et al. 2010 for an example with qPCR data in rodent malaria infections). Ideal methods for estimating population size would produce consistent coefficients of variation whether individuals are abundant or rare (e.g., negative binomial error distribution). For the case of malaria infections, our new approach allows for sampling schedules that are entirely feasible in experimental animal models (Deharo et al., 1994, 1996; O’Donnell et al., 2011; Prior et al., 2018; O’Donnell et al., 2022) and even controlled human infection trials (reviewed in Duncan and Draper 2012). For plants and animals, a wealth of data exist that can be used to parameterize matrix models (Salguero-Gómez et al., 2015, 2016), though the goal is typically to estimate population expansion rates from age distribution data (Hernández et al., 2023) rather than the reverse.

The methods in the present study rely on time series of abundance, and there are important challenges to obtaining accurate abundance data. In particular, the question of whether synchrony can be resolved from the data depends on the level of sampling noise, which for malaria parasites requires replicate samples from the same infection at the same point in time. These data exist for PCR-derived counts in *P. chabaudi* (Mideo et al., 2008; Huijben et al., 2010; Miller et al., 2010), but comparable data from microscopy-derived iRBC counts appear to be lacking in both *P. chabaudi* and *P. falciparum*. Estimates of that sampling error are needed to make specific recommendations about experimental design, including sampling frequency and duration. All else equal, per capita replication rates will approach one (the population replacement level) as sampling resolution increases, because increases in iRBC abundance will be subdivided into smaller and smaller intervals. That could make it more difficult to distinguish synchrony-driven differences in per capita replication rates and may represent another reason to aim for clustering sampling around median bursting times, when changes in iRBC abundance—and therefore in *R_e_* values—will be the greatest. Further, if error distributions were known, it would be possible to implement a maximum likelihood approach rather than minimizing the sum absolute error as we have done here.

The other major challenge is that some developmental ages are more difficult to sample. For example, the mature stages of some *Plasmodium spp.* sequester where they cannot be sampled, at least in certain hosts. When sequestration occurs, it would lead to underestimates of total parasite abundance (e.g., *P. falciparum*, White et al. 1992; *P. vivax*, Carvalho et al. 2010; *P. berghei* in wild type but not *rag1-/-* mice, Khoury et al. 2014). If the level of synchrony is changing through time, the bias introduced by sequestration may likewise fluctuate, making it difficult to compare synchrony estimates from different phases of infection or across different infections. Sequestration is not universal, so some host-parasite combinations would not suffer from this bias (including *P. berghei* in *rag1^−/−^* mice, Khoury et al. 2014, any *P. falciparum* infections *in vitro*). Sequestration is also a problem for studies attempting to quantify synchrony *in vivo* from age distribution data (Deharo et al., 1994, 1996; Touré-Ndouo et al., 2009; O’Donnell et al., 2011; Ciuffreda et al., 2020), along with the added challenge of the bias introduced by population dynamics (reviewed in Greischar et al. 2019). Accounting for the bias introduced by sequestration will be crucial for making comparisons across parasite species that also vary in their propensity to sequester as well as across environments where the potential for sequestration varies (e.g., *in vivo* and *in vitro* conditions).

Empirical advances offer some promise for sidestepping the problem of sequestration to estimate total iRBC abundance from proxy measures. Methods exist to quantify the total number of parasite genomes from volatiles in breath samples (Berna et al., 2015), plasma biomarkers (Dondorp et al., 2005), and—for experimental animal models—from transgenic parasite strains expressing luciferase or other markers (Franke-Fayard et al., 2005). Caution is warranted in using these methods, since they quantify parasite genomes, rather than iRBC abundance. Unfortunately, those two quantities are only equivalent for parasite populations that are synchronized and have not yet begun DNA replication (e.g., Cheesman et al. 2003), so studies attempting to quantify total parasite biomass must make assumptions regarding the synchrony of the parasite population being examined. The present study suggests that if biomarker abundance were quantified at multiple time points per IDC, it may be possible to fit rather than assume the underlying level of synchrony by modifying the approach introduced here. As with microscopy-estimated counts, quantifying the sampling error distribution associated with these newer methods will be crucial to determining feasibility.

Perhaps even more challenging is the fact that time series of iRBC abundance are not typically gathered when the goal is estimating synchrony; rather, stage percentages are overwhelmingly used to compare synchrony through time and across different infections (reviewed in Greischar et al. 2019). The focus on stage percentages is understandable, given that researchers often require parasite populations enriched for a particular developmental age class (for example, when testing stage-specificity of antimalarial drugs, e.g., ter Kuile et al. 1993). However, knowing the stage percentages at a particular point in time does not enable the projection of future stage percentages, especially when parasite numbers are changing (Greischar et al., 2019). Since it utilizes replication rates, our new approach can project how age distributions will change through time for a given initial level of synchrony. Projected age distributions could prove useful, for example, in predicting when parasite populations will be most vulnerable to drug treatment or immune clearance.

Existing methods for quantifying synchrony are useful when abundance is steady through time and the duration of morphologically-distinguishable developmental stages is known in advance (reviewed in Greischar et al. 2019). Those conditions are likely to hold for only a small minority of cases, but the method validated here can estimate synchrony when numbers are changing—and in fact works best when populations are undergoing dramatic changes in abundance—and requires no prior knowledge of stage durations. The key requirements of this approach are time series of abundance, sampled multiple times per developmental cycle, and knowledge about the duration of development. While the focus with stage percentage data has left a dearth of such data, they are obtainable (O’Donnell et al., 2022). Further, given the kind and resolution of data required, the approach is likely to have utility in other organisms, both pathogenic and free-living. Past studies quantifying changes in synchrony in free living organisms have relied on time series of abundance (e.g., insect pests, Yamanaka et al. 2012; Nelson et al. 2013; Bjørnstad et al. 2016), but we have shown that a useful signal of synchrony also emerges from per capita replication rates. Whether models fit abundance or per capita growth rates, accounting for population dynamics is necessary to identify ecological processes that maintain versus erode synchrony and to determine whether synchrony is heritable and capable of evolving. These unresolved questions carry enormous practical significance—the answers can inform the schedule of drug treatment (or for free-living pests, chemical controls) and enable predictions about evolutionary responses to intervention efforts. For malaria parasites, better understanding of variation in synchrony—which requires robust methods to estimate it—could improve understanding of the pathogenesis and epidemiology and inform efforts to intervene at the individual and population level.

## AUTHOR CONTRIBUTIONS

Megan A. Greischar: conceptualization, investigation, methodology, formal analysis, funding acquisition, and writing–original draft, validation, visualization; Nicholas J. Savill: conceptualization, funding acquisition, writing–review and editing, visualization; Sarah E. Reece: conceptualization, funding acquisition, writing–review and editing, visualization; Nicole Mideo: conceptualization, funding acquisition, project administration, writing–review and editing, validation, visualization. All authors contributed to drafts and gave final approval for publication.

## Supporting information

Supplemental code and simulated data

## ACKNOWLEDGMENTS

We have benefited greatly from discussions with Stephen Ellner, Christina Hernández, Timothy Lambert, Aidan O’Donnell, Aĺız Owolabi, Damie Pak and Jordi Ripoll.

## DATA AVAILABILITY STATEMENT

This study uses simulated data, and all code needed to reproduce the analysis and figures is annotated and available in supplemental files.

## SUPPLEMENTAL FIGURES

**Figure S1.**
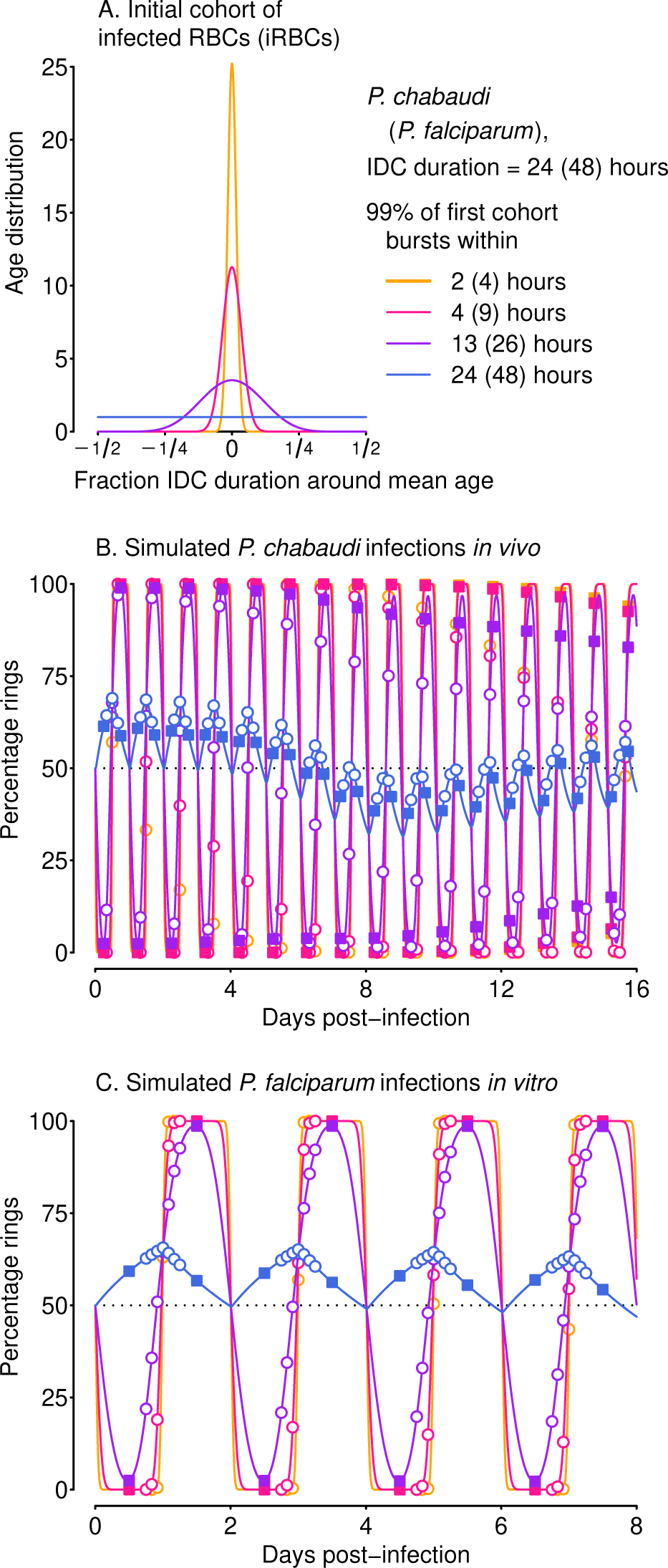
Percentage rings for the simulations shown in fig. 1. The bursting for the initial cohort is shown in panel A (identical to fig. 1A), with the percentage rings in *P. chabaudi* and *P. falciparum* shown in panels B and C, respectively. Closed squares (open circles) indicate time points corresponding to even (uneven) sampling. See main text for details.

**Figure S2.**
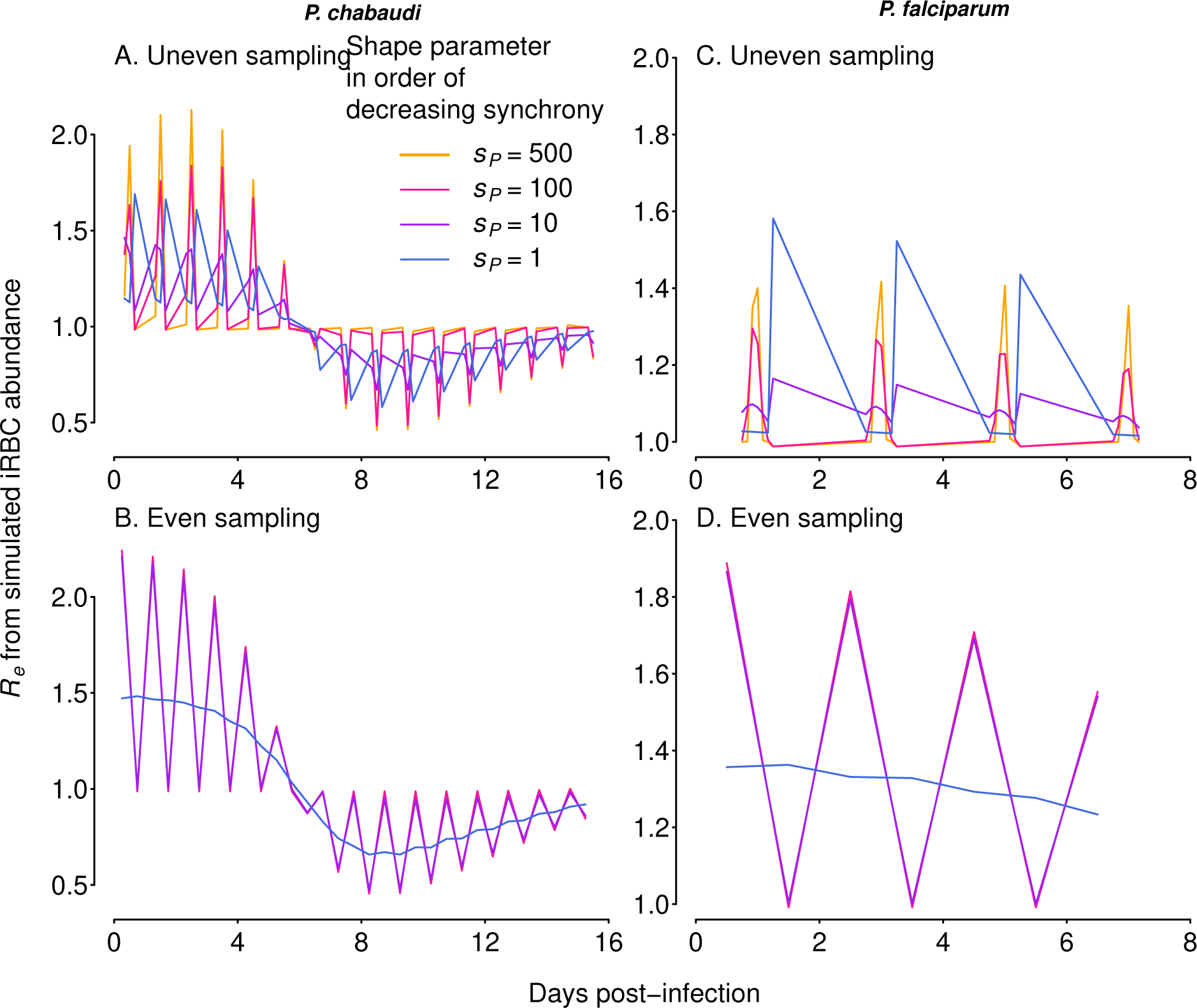
Simulated per capita replication rates (*R_e_*) can show differences in synchrony more clearly than iRBC abundance (compare to figure 1B, C). Left panels show *R_e_* values for simulated *P. chabaudi* (A) and *P. falciparum in vitro* (B) infections sampled unevenly, while right panels show *R_e_* values when infections are sampled twice per IDC (C, D). Note that *R_e_* values vary with sampling schedule, and sparse sampling (panels C, D) can make it difficult to discern smaller differences in synchrony.

**Figure S3.**
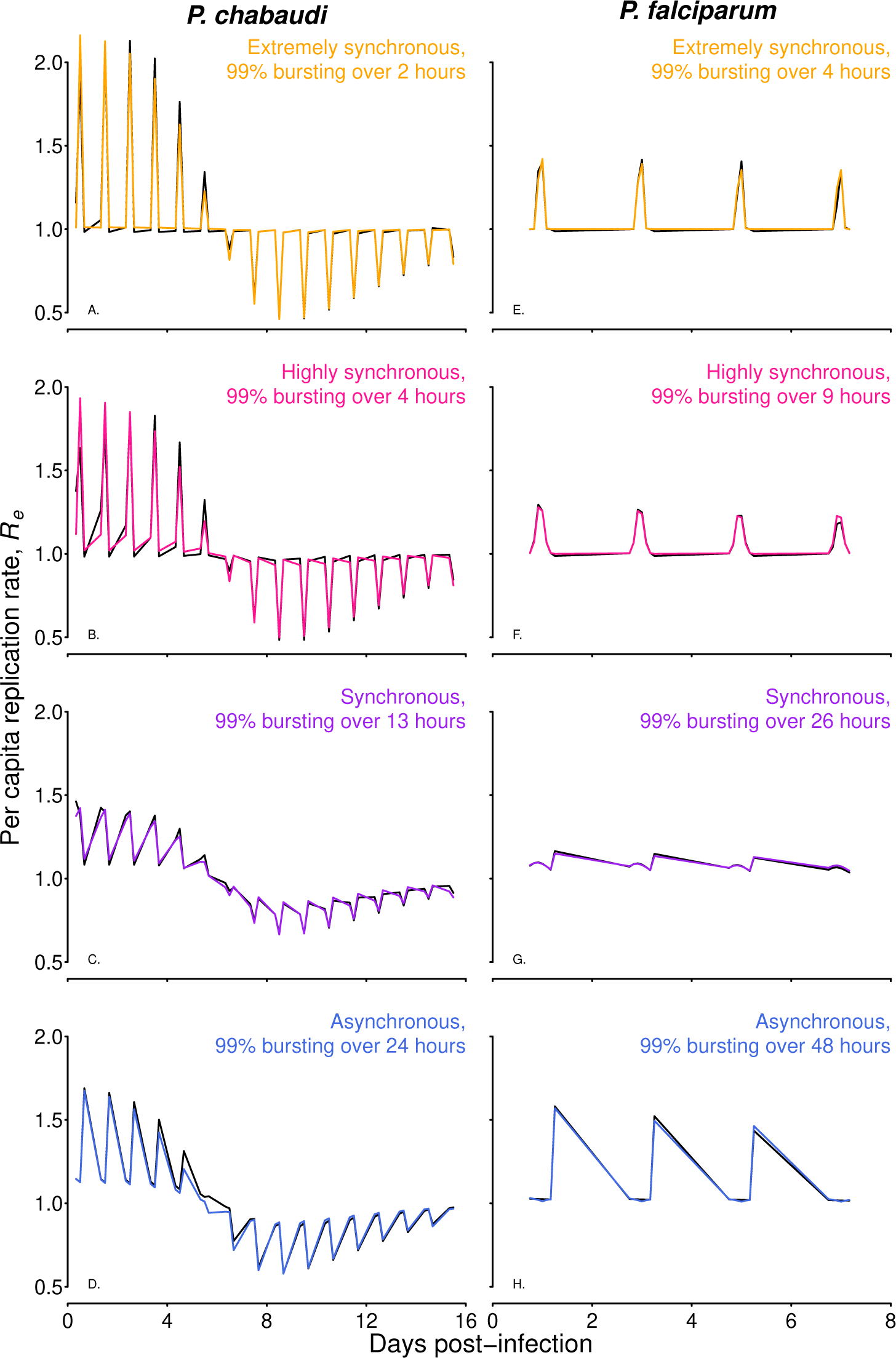
The fitted model can closely track simulated time series sampled at uneven intervals. True (that is, derived from simulated “observations”, black) and fitted per capita replication rates (Re and ^Re, respectively) are shown for simulated P. chabaudi (left panels) and P. falciparum (right panels) time series. Colors as in figure 1, and sampling occurred 3-4 times daily for each species (see Simulating experimental infection data section in Methods for details). The Re values shown in black are identical to the trajectories overlaid in figure S2A, C.

**Figure S4.**
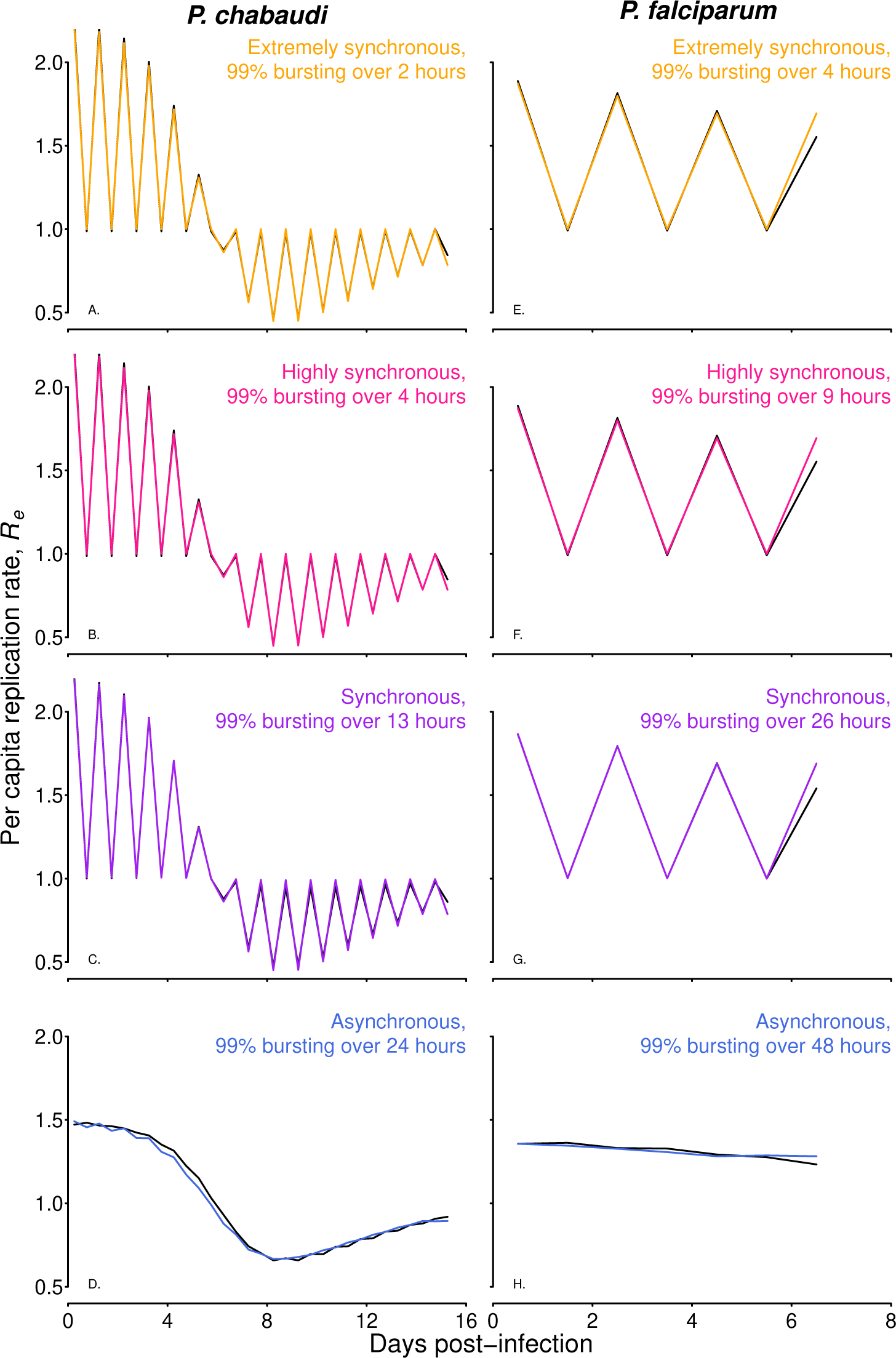
The fitted model can closely track simulated time series sampled at even intervals. True (derived from simulated “observations”, black) and fitted per capita replication rates (Re and ^Re, respectively) are shown for simulated P. chabaudi (left panels) and P. falciparum (right panels) time series. Colors as in figure 1, with sampling occurring at even intervals, twice per IDC. The Re values shown in black are identical to the trajectories overlaid in figure S2B, D.

**Figure S5.**
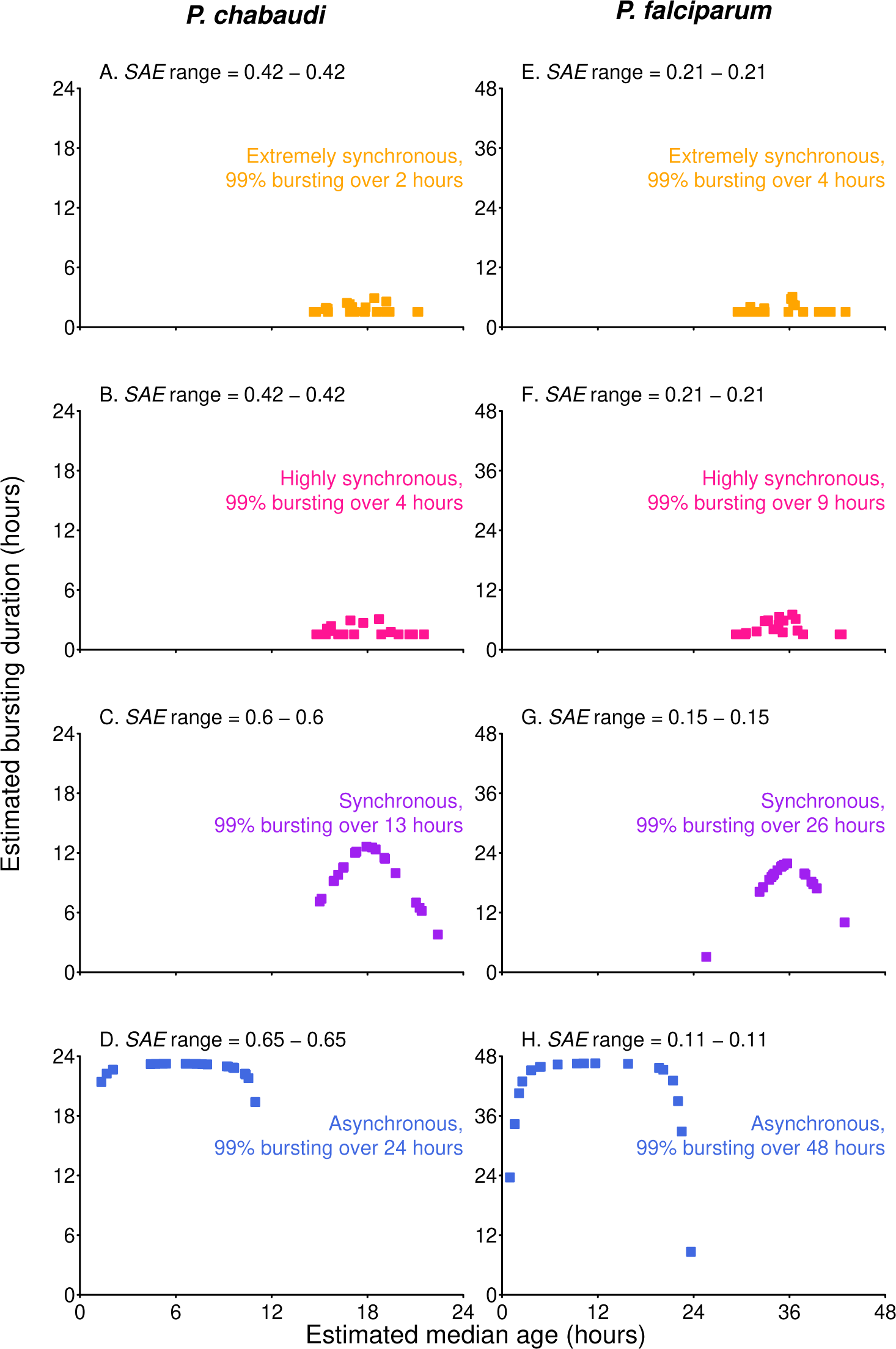
With even sampling, multiple combinations of median parasite age and duration of bursting yield identical fits (nonidentifiability). For both species and each level of synchrony simulated, the combinations of best fitting median parasite age and bursting duration are shown, with the range of sum absolute error (SAE) values for even sampling given in the plot title. Open circles indicate corresponding fits for uneven sampling, for reference (points lie on top of each other). Colors as in figure 5.

**Figure S6.**
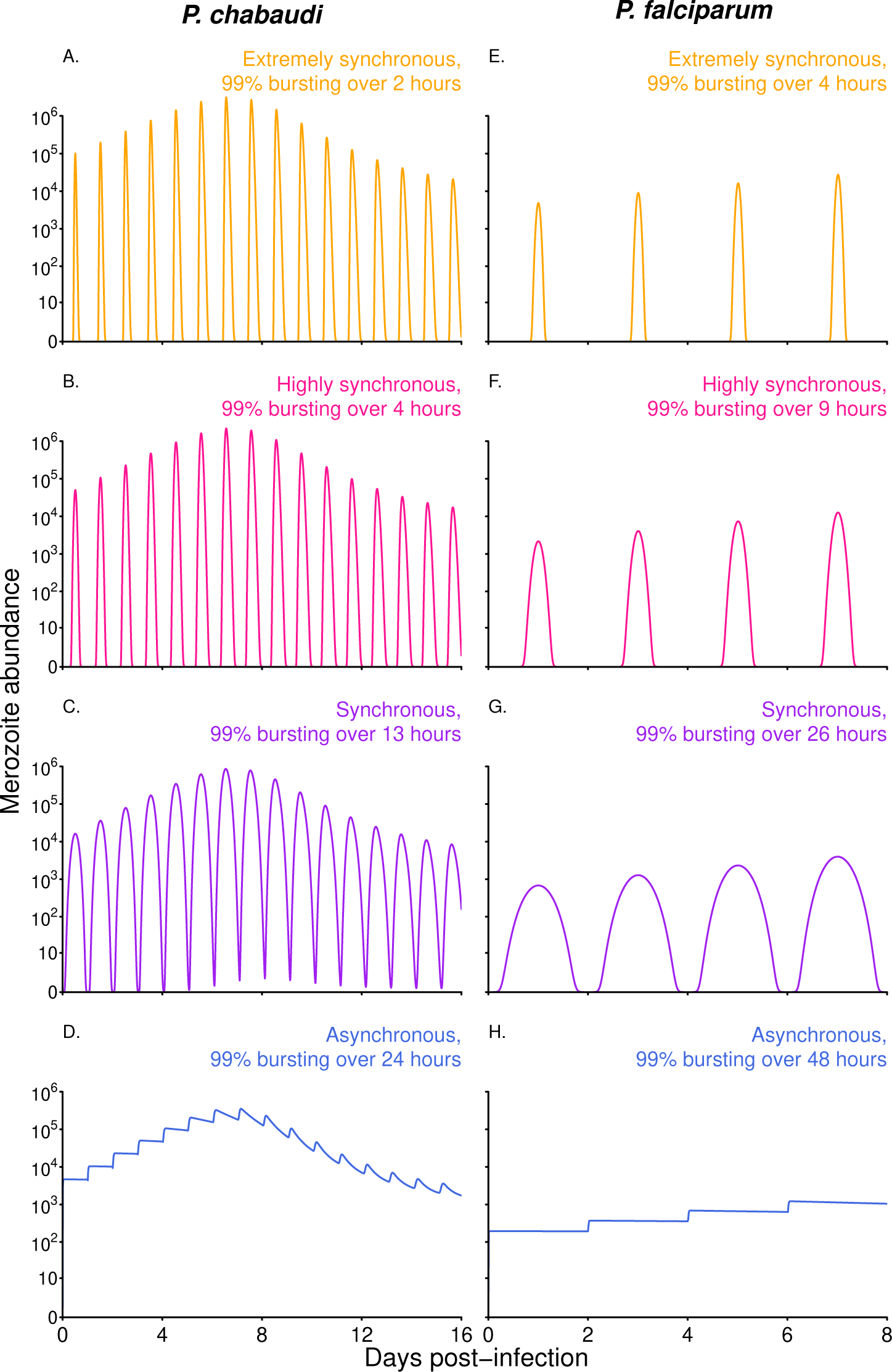
The duration of bursting is apparent from merozoite abundance in the simulated infections shown in fig. 1. Simulated *P. chabaudi* infections encompass 16 IDCs (A-D), and synchrony decays to a small but noticeable degree in extremely and highly synchronous infections (note the widening of peaks over the course of infection). In contrast, simulated *P. falciparum* infections (E-H) encompass only 4 IDCs and synchrony has minimal opportunity to decay.

**Figure S7.**
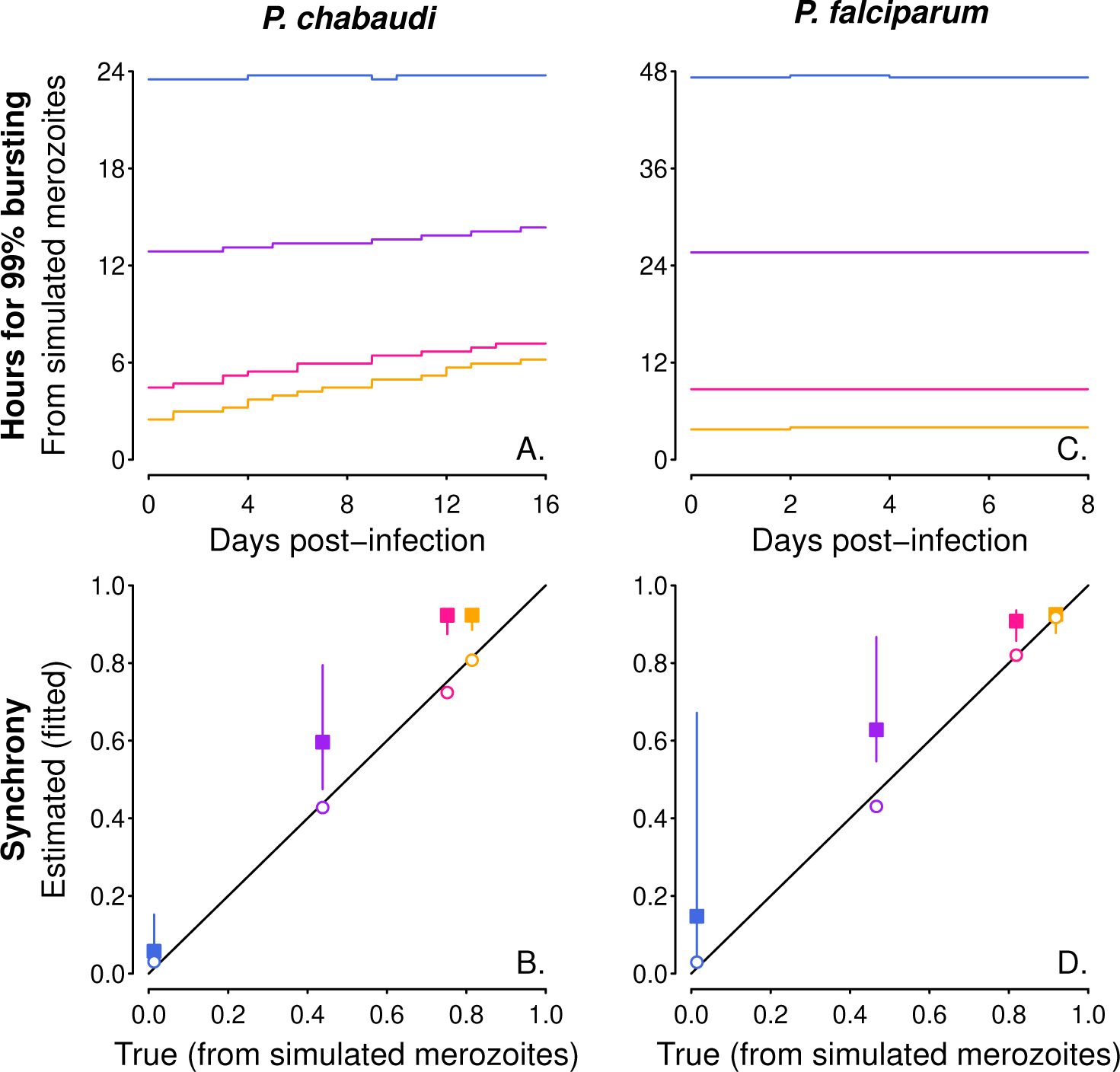
Changes in synchrony through time (or a lack thereof) are apparent when bursting duration is estimated from simulated merozoite abundance, and the new approach can estimate the correct average level of synchrony with uneven sampling. The most apparent changes in synchrony are in the extremely and highly synchronized simulated infections for *P. chabaudi*, which encompasses 16 IDCs (A), while *P. falciparum* infections show minimal changes in synchrony over the 4 IDCs simulated (B). Calculating synchrony using Eq. 1 from the average duration of bursting shown in panels A and B yields the x-axis values in panels C and D (respectively). When changes in bursting duration through time are incorporated, the new method returns estimates nearly identical to the true values (C, D) for uneven sampling (open circles). As in figure 5, even sampling (closed squares) results in non-identifiability such that a range of estimated synchrony values give similar fits to data (see figure S5).

**Figure S8.**
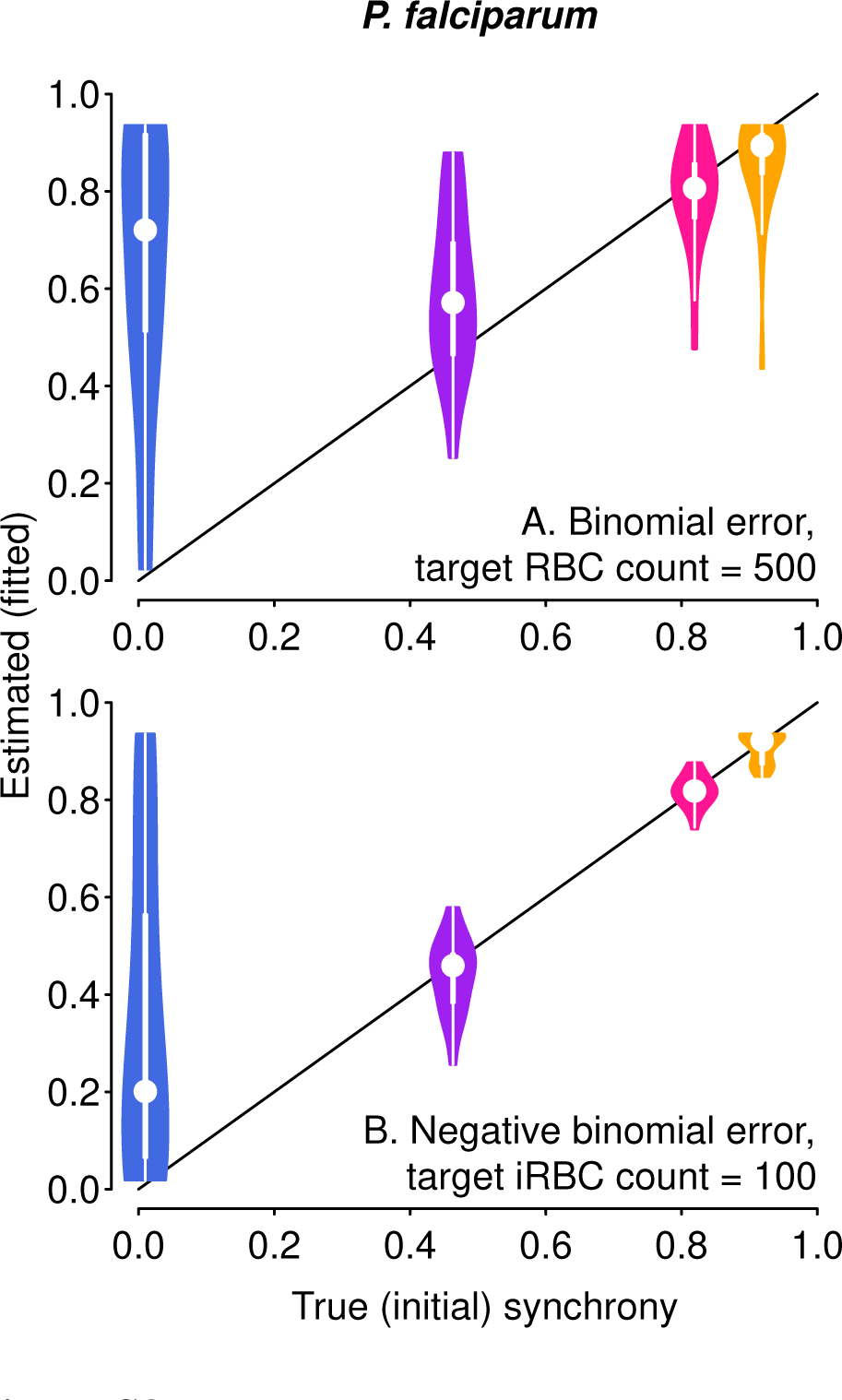
Synchrony differences are harder to discern for *P. falciparum*, which was simulated to maintain a low percent parasitemia (approx. 10% or less, Table S2). Target counts as in fig. 6A, C.

**Figure S9.**
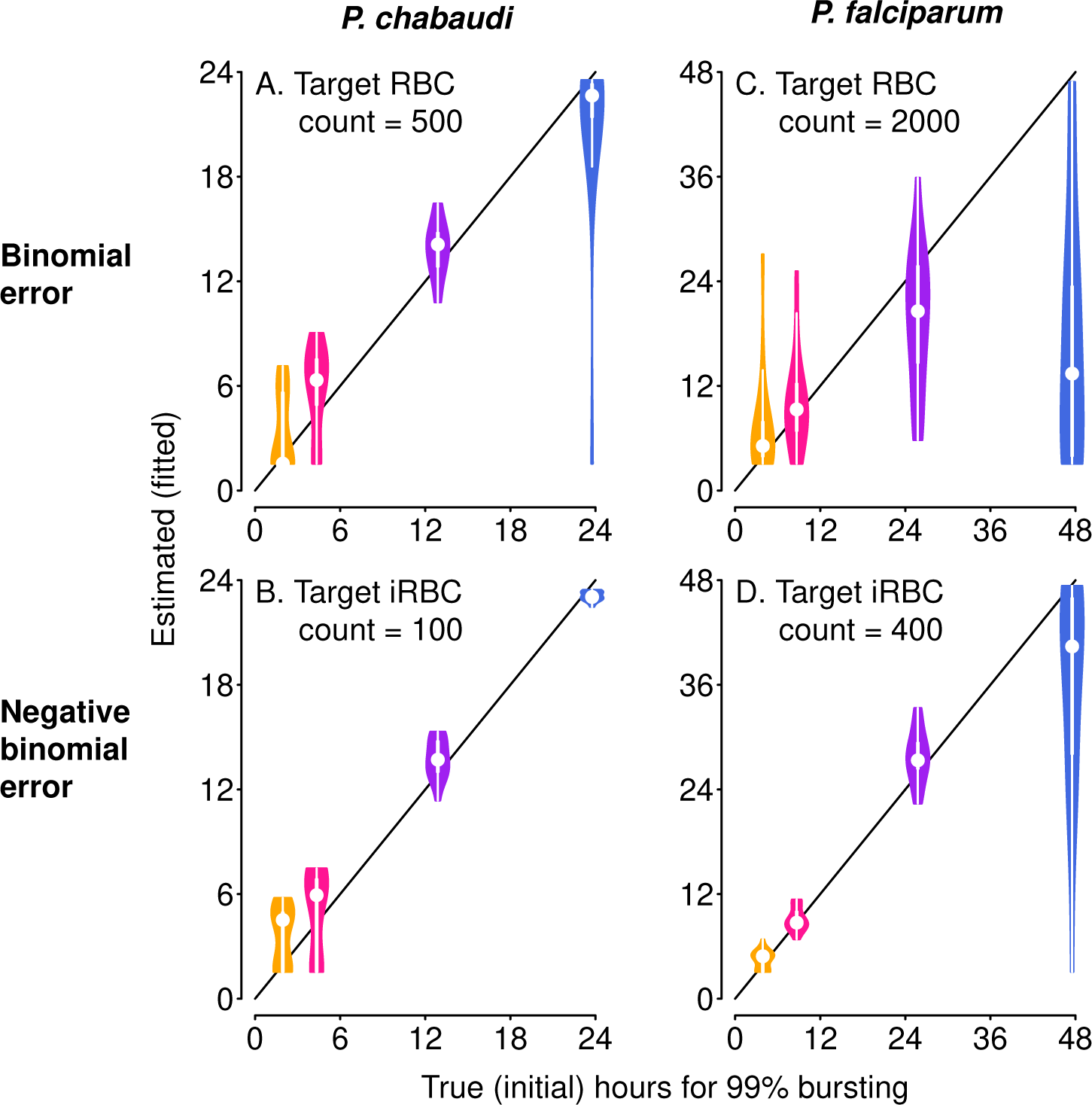
The ability to distinguish differences in the duration of bursting depends on distribution of sampling noise. Violin plots showing the distribution of 20 fits to time series with simulated sampling noise, with a binomial (A, B) or negative binomial (C, D) distribution. When error is binomially distributed (A, B), differences are difficult or impossible to detect, with asynchronous infections often appearing with a spuriously short duration of bursting (that is, highly synchronized). Duration of bursting estimates are closer to true values when sampling noise follows a negative binomial distribution (C, D).

## SUPPLEMENTAL METHODS

### Simulating experimental infection data

Whether the focus is *in vivo* or *in vitro* infections, iRBC abundance (defined in Table 1) can be readily obtained in experiments using a cell counter to estimate total RBC counts and microscopy of blood smears to estimate the percentage of RBCs that are infected (e.g., Taylor et al. 1997). Many existing studies focus on parasite population dynamics rather than synchrony, and so report iRBC abundance once per IDC, while other studies examine synchrony and report stage percentages for multiple time points during each IDC but without the corresponding total RBC counts (Lambros and Vanderberg, 1979; Deharo et al., 1994, 1996; Reilly et al., 2007; Touré-Ndouo et al., 2009; Allen and Kirk, 2010; O’Donnell et al., 2011). What is needed for our approach is iRBC abundance, sampled at multiple times per IDC; such data have been obtained for both *P. chabaudi* (Prior et al., 2018; O’Donnell et al., 2022) and *P. falciparum* infections *in vivo* (including from historical human data, e.g., Eichner et al. 2001).

We simulated malaria infections of mice (*P. chabaudi*) and *in vitro* cultures of human malaria parasites (*P. falciparum*), tracking abundance of iRBCs. The model comprises a set of delayed differential equations, where the delay *α* specifies the length of the IDC and is assumed to have a fixed duration (Greischar et al., 2014), where *α* = 1 day (24 hours) for *P. chabaudi*, and *α* = 2 days (48 hours) for *P. falciparum*. Merozoites (the RBC-invasive forms) are assumed to lose viability at a constant rate, yielding an exponential distribution of lifespans in the absence of RBCs to invade. Merozoites remain viable for only minutes on average (see Tables S1, S2 for all parameter values) and are lost rapidly from circulation when there are RBCs available to invade. Therefore, even though the merozoite stage is not fixed in duration, it is so brief that the level of synchrony imposed by the initial age distribution of parasites tends to be maintained through time (Greischar et al., 2014). We considered four scenarios with simulated time series of iRBC abundance, ranging from an extremely high initial level of synchronization to complete asynchrony (figure 1). The initial level of synchrony is defined by the shape parameter of the beta distribution, *s_P_* (Table 1), which can be used to specify the length of time required for 99% of the initial cohort to burst (figure 1A). For simplicity, simulated infections lack any host immune response or transmission stage production. Still, the model outputs capture the key population dynamic features that undermine existing methods: iRBC numbers increase several orders of magnitude, and for the longer *P. chabaudi* simulations decrease over the same scale (due to red blood cell limitation; figure 1B, C).

To simulate experimental data, we subsampled iRBC abundance from the malaria infection models from time points that mimic plausible experimental designs. For *P. chabaudi* simulations, iRBC abundance was sampled at three uneven intervals per developmental cycle, since sampling more frequently from individual hosts is likely to be infeasible. We assumed that sampling was centred around peak bursting. Specifically, the peak time of initial bursting occurs at *t* = IDC*/*2 (i.e., at *t* = 1*/*2 day for *P. chabaudi*, corresponding to zero in figure 1A). Subsequently, peak bursting clusters around *t* = 1*/*2 + *d*, where *d* indicates the day of sampling, with some small variation due to the duration of the merozoite stage. We sample our simulated infections in an eight-hour window beginning four hours before bursting, so that iRBC abundance is obtained at *t* = 1*/*2 + *d* − 4*/*24, *t* = 1*/*2 + *d*, and *t* = 1*/*2 + *d* + 4*/*24 for each day. For *Plasmodium falciparum*, we are simulating in vitro infections, where more samples are possible. Here, we sample seven times, every two-hours for the 12-hour period spanning peak bursting, as described by (Reilly Ayala et al., 2010). Sampling more frequently is likely to be infeasible, especially for *in vivo* infections, and we also consider lower resolution time series, where iRBC populations are sampled at even intervals twice per IDC, either six hours before and after peak bursting (*P. chabaudi*), or 12 hours before and after peak bursting (*P. falciparum*).

### Adding sampling noise to iRBC count data

Quantifying iRBC abundance requires a total RBC count and an estimate of the fraction of RBCs occupied by parasites (percent parasitemia), and each of these two steps can contribute to sampling noise. The sampling error inherent to estimating total RBC abundance with a cell counter has been estimated to be normally distributed around log_10_-transformed RBC counts with a standard deviation of 0.034 (Mideo et al., 2008), so we add random noise from that distribution to our deterministic simulations to mimic replicate mice in an experiment. For simplicity, we assume that total RBC counts obtained for *P. falciparum* infections *in vitro* follow the same error structure. The error distribution of percent parasitemia estimates is less clear, so we simulate two error distributions—binomial and negative binomial—that might be expected to emerge from microscopy protocols. *P. falciparum* infections are typically assessed by counting a certain number of RBCs regardless of infection status (either 500 or 2000 RBCs, Division of Parasitic Diseases and Malaria 2016). We assume that process generates binomially-distributed errors, in which the coefficient of variation declines as parasitemia increases, as has been reported for parasitemia estimated by microscopy in *P. falciparum* (O’Meara et al., 2005). For *P. chabaudi* infections, a common protocol is to count infected and uninfected RBCs until a target number of iRBCs is achieved, either 100 (Chimanuka et al., 1999) or 400 (P. Schneider, *pers. comm.*). That process may be expected to yield a negative binomial distribution of error, and a relatively consistent coefficient of variation in percent parasitemia estimates whether the true parasitemia is low or high. For context, *P. chabaudi* infections can attain 70% parasitemia (Chimanuka et al., 1997) while *P. falciparum* cultures are typically maintained so as to keep parasitemia below 5% (Reilly et al., 2007).

**Table S1.** Parameters for within-host models of *P. chabaudi* infections of mice. Initial inoculum size was chosen to give a reasonable approximation of the dynamics of many experimental *P. chabaudi* infections, which attain peak parasite density at day seven post-infection, with RBCs subsequently dropping to their minimum value, on the order of a million RBCs per *µ*L (Huijben et al., 2010). Note that we used a larger merozoite mortality rate (*µ_z_*) than that used previously (Greischar et al., 2014, 2016) with the aim of slowing the loss of synchrony.

| Parameter | Value | Source |
| --- | --- | --- |
| Cycle length, $\alpha$ | 1 day | Landau and Boulard (1978) |
| Merozoite mortality rate, $\mu_z$ | 72/day | assumed by Gravenor et al. (1995) |
| Burst size, $\beta$ | 10 merozoites | Mideo et al. (2008) |
| Initial inoculum size, $I(0)$ | $\approx 44000$ per $\mu\text{L}$ | see caption |
| Invasion rate, $p$ | $2.5 \times 10^{-6} \text{ day}^{-1} \text{ merozoite}^{-1}$ | Mideo et al. (2008) |
| RBCs per $\mu\text{L}$, $R(0)$ | $8.5 \times 10^6$ | Savill et al. (2009) |
| RBC mortality rate, $\mu$ | 0.025/day | Miller et al. (2010) |
| Max. rate of erythropoiesis, $\lambda$ | $3.7 \times 10^5 \text{ RBCs}/\mu\text{L}/\text{day}$ | Savill et al. (2009) |

**Table S2.** Parameters for within-host models of *P. falciparum* infections *in vitro*. Invasion rate and initial inoculum size were chosen so that the percentage iRBCs expanded from 1% to 10.7% over the 8 days simulated.

| Parameter | Value | Source |
| --- | --- | --- |
| Cycle length, $\alpha$ | 2 days | Garnham (1966) |
| Merozoite mortality rate, $\mu_z$ | 200/day | Boyle et al. (2010) |
| Burst size, $\beta$ | 16 merozoites | Reilly et al. (2007) |
| Initial inoculum size, $I(0)$ | $0.01 \times [R(0) + I(0)]$ | see caption |
| Invasion rate, $p$ | $5 \times 10^{-5}$ day<sup>-1</sup> per merozoite | see caption |
| Initial RBCs per $\mu\text{L}$, $R(0) + I(0)$ | 555555.6 | Loe and Edwards (2004) report $5 \times 10^6$ RBCs/$\mu\text{L}$ and 45% hematocrit is within a realistic range for humans (reviewed in Dean 2005). From these values, we calculate the number of cells in 5% hematocrit, the range desirable for *P. falciparum* *in vitro* (e.g., Allen and Kirk 2010), by multiplying $5 \times 10^6$ by 0.05 and dividing by 0.45. |
| RBC mortality rate, $\mu$ | 1/120 days | Koury and Ponka (2004) |
| Max. rate of erythropoiesis, $\lambda$ | 0 | no RBC replenishment *in vitro* |

